# Local unfolding of the HSP27 monomer regulates chaperone activity

**DOI:** 10.1101/345751

**Authors:** T. Reid Alderson, Julien Roche, Heidi Y. Gastall, Iva Pritišanac, Jinfa Ying, Ad Bax, Justin L. P. Benesch, Andrew J. Baldwin

## Abstract

The small heat-shock protein HSP27 is a redox-sensitive molecular chaperone that is expressed throughout the human body. Here we describe redox-induced changes to the structure, dynamics, and function of HSP27 and its conserved α-crystallin domain, and provide the first structural characterization of a small heat-shock protein monomer. While HSP27 assembles into oligomers, we show that the transiently populated monomers released upon reduction are highly active chaperones *in vitro*, but are kinetically unstable and susceptible to uncontrolled aggregation. By using relaxation dispersion and high-pressure nuclear magnetic resonance spectroscopy, we reveal that the pair of β-strands that mediate dimerization become partially disordered in the monomer. Strikingly, we note that numerous HSP27 mutations associated with inherited neuropathies cluster to this unstructured region. The high degree of sequence conservation in the α-crystallin domain amongst mammalian sHSPs suggests that partially unfolded monomers may be a general, functional feature of these molecular chaperones.

## Introduction

Small heat-shock proteins (sHSPs) are a class of molecular chaperones present in all kingdoms of life and exhibit diverse functionality, from modulating protein aggregation to maintaining cytoskeletal integrity and regulating apoptosis^1^. Despite molecular masses in the range of 10-40 kDa, sHSPs assemble into large, heterogeneous oligomers^2^ whose constituent monomers and dimers typically exchange between oligomers^3^. The chaperone activities of many sHSPs have been characterised *in vitro*^4^, but the active sHSP species remains unclear, with large oligomers^5,6^, small oligomers^7,8^, and dimers^9^ all implicated.

The most abundant sHSP in humans, HSP27 (or HSPB1), is systemically expressed under basal conditions and upregulated by oxidative stress^10^, during aging^11^, and in cancers^12^ and protein deposition diseases^13^. Numerous mutations in HSP27 have been linked to different neuropathies, including distal hereditary motor neuropathy (dHMN) and Charcot-Marie-Tooth (CMT) disease^14,15^, the most commonly inherited neuromuscular disorder. These maladies are themselves linked to oxidative stress^16,17^, and recent studies have indicated that the reducing environment of the cytosol progressively transitions to an oxidising environment over the lifetime of an organism^18,19^.

HSP27 is directly sensitive to the intracellular redox state via its lone cysteine residue (C137), which controls dimerization by forming an intermolecular disulphide bond *in vivo*^20^. This cysteine is highly conserved in HSP27 orthologs but not found in other mammalian sHSPs^21^, implying that it plays an important and specific role in regulating function. Accordingly, the presence of this disulphide bond impacts on the activity of HSP27 *in vitro*^22–24^, and on the resistance of cells to oxidative stress^20,21,25,26^. Intriguingly, variants of HSP27 that have an increased tendency to form monomers display hyperactivity both *in vitro* and *in vivo*^27,28^. More generally, sHSP monomers can mediate the subunit exchange between the oligomeric assemblies^3,29^. However, because they are typically present at low abundance in solution, no sHSP monomer has yet been characterised structurally.

Obtaining high-resolution information describing HSP27 is challenging, as it assembles into a polydisperse ensemble of inter-converting oligomers ranging from approximately 12 to 36 subunits^30–32^ of average molecular mass of ca. 500 kDa. However, removal of the C-terminal region (CTR) and N-terminal domain (NTD) leaves a conserved ~80-residue, ‘α-crystallin’ domain (ACD) that does not assemble beyond a dimer (Fig. 1a). The subunits in the dimer adopt an immunoglobulin-like fold, and assemble through the formation of an extended β-sheet upon pairwise association of their β6+7 strands^33–36^. Under oxidising conditions, the dimer interface in HSP27 is reinforced by an intermolecular disulphide bond involving C137 from adjacent subunits centred on a two-fold axis^33–35^. Based on evidence from the closely related sHSP, αB-crystallin^37^, the ACD is likely structurally similar in the context of the full-length oligomeric protein and in its isolated dimeric form. Moreover, as the excised ACD of αB-crystallin has been shown to be incorporated into full-length sHSP oligomers^38^ and display potent chaperone activity *in vitro*^34,39^, it appears that important aspects of sHSP function are encoded within this domain.

Here, we have employed an integrative biophysical approach to interrogate the impact of redox-induced changes to the structural features of HSP27 and its excised ACD (cHSP27). We find that reduced HSP27 more effectively prevents protein aggregation than its oxidised counterpart. However, neither the distribution of HSP27 oligomers, nor the conformations and fast dynamics of the CTR and cHSP27 dimer vary appreciably with oxidation state. Rather, upon cleavage of the disulphide bond, the release of monomers into solution is responsible for the observed differences in chaperone activity upon reduction. To interrogate the structure of the free, but sparsely populated, monomers we have used a combination of Carr-Purcell-Meiboom-Gill (CPMG) relaxation dispersion (RD) and high-pressure solution-state nuclear magnetic resonance (NMR) spectroscopy methods. Our data reveal that monomeric cHSP27 becomes highly dynamic and disordered in the region that previously constituted the dimer interface. While we find the monomer to be highly chaperone-active *in vitro*, we demonstrate that increasing the abundance of the monomer results in a heightened tendency for uncontrolled self-aggregation. The importance of the unstructured region in this delicate balance between function and malfunction can be linked by mutations in HSP27 that are associated with hereditary neuropathies, which mainly cluster to the disordered region of the monomer.

## Results

### Reducing HSP27 affects chaperone activity by increasing the amount of free monomer

We first examined full-length HSP27 to analyse redox-dependent changes to its oligomeric distribution. Native mass spectra of reduced and oxidised HSP27 were highly similar, with overlapping signals in the 5,000–15,000 m/z region (Fig. 1a), consistent with previous data^30,31^ This reveals that HSP27 assembles into large, polydisperse oligomers with similar distributions under both conditions (Supplementary Fig. 1). We also observed monomeric and dimeric HSP27 in the spectra of both oxidised and reduced forms, with a significant increase in the population of free monomer upon reduction (Fig. 1a).

To confirm that dissociation of the dimers, rather than modulation of the oligomers, is the major consequence of reduction, we used solution-state NMR to examine the CTR, which can mediate the assembly of sHSPs^40,41^. As HSP27 oligomers have an average mass of ca. 500 kDa, only the disordered CTR from ^15^N labelled HSP27 can be observed in a 2D ^1^H-^15^N heteronuclear single quantum coherence (HSQC) NMR spectrum^42–44^. To probe the local dynamics in this region quantitatively, we recorded NMR spin relaxation experiments that characterize motions on the ps-ns timescale (Supplementary Fig. 1). No significant differences in the conformation or fast backbone motions were detected between oxidised and reduced forms of HSP27. Our combined native MS and NMR data on the polydisperse ensemble populated by HSP27 demonstrate that the primary impact of reduction is the release of free monomers.

To ascertain whether the presence of monomers impacts on chaperone function, we used the model substrate citrate synthase (CS)^4,45^ to probe the activity of HSP27 *in vitro*. The aggregation of CS alone was suppressed upon the addition of reduced HSP27 at low concentrations (0.5 μM) (Fig. 1b). The reduced sample contains a mixture of monomers and dimers, whereas the in the oxidised sample the sub-oligomeric species are predominantly dimeric^31^. These data therefore suggest that monomerisation regulates the chaperone activity of HSP27, rendering it more effective at suppressing aggregation *in vitro*.

**Fig. 1:**
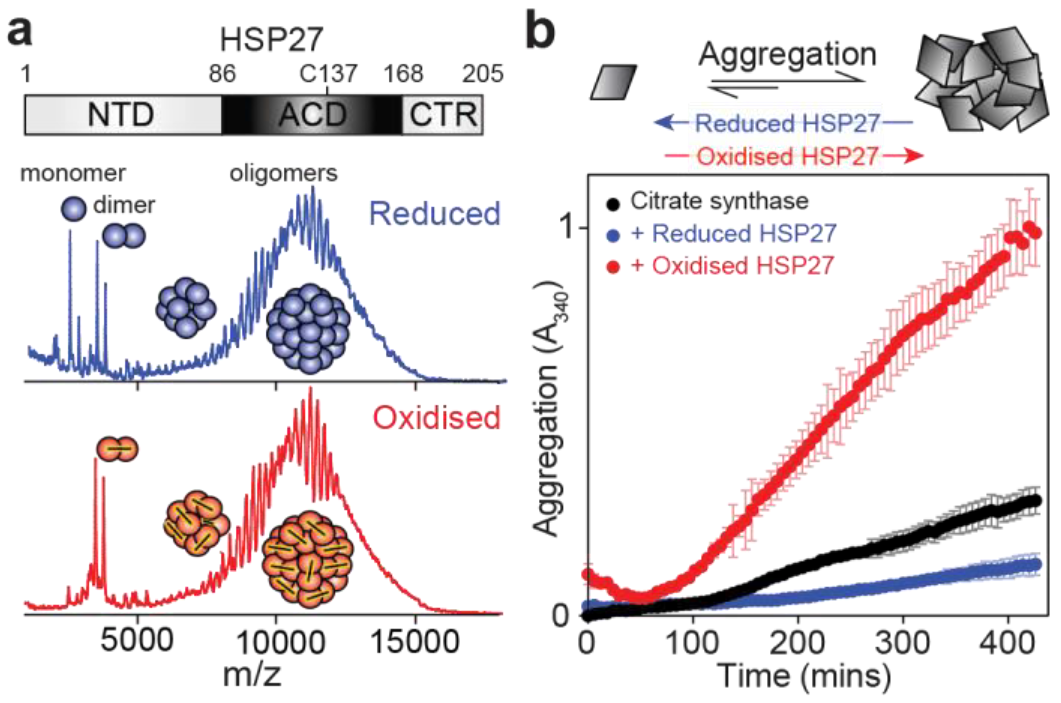
Reduction of HSP27 releases monomers from the polydisperse oligomers. (**a**) Domain architecture of the human molecular chaperone HSP27, which forms polydisperse oligomers that reach >500 kDa. Native mass spectra collected at 25 μM total monomer concentration for both oxidised (*red*) and reduced (*blue*) HSP27 reveal the formation of polydisperse oligomers. More monomers are present in the reduced sample, but the oligomeric distributions are highly similar (Supplementary Fig. 1). (**b**) The chaperone activity of HSP27 was assayed by monitoring the increase in light scattering at 340 nm of 10 μM of a thermo-sensitive substrate (CS) in the presence or absence of 0.5 μM reduced (*blue*, 5 mM 2-mercaptoethanol, BME) or oxidised (*red*) HSP27. The average and ± standard deviation of two replicates are indicated.

### *HSP27 monomers are potent chaperones* in vitro *that readily self-aggregate*

To examine these redox effects on the monomer:dimer equilibrium in the absence of oligomeric forms of HSP27, we turned to the truncated form, cHSP27 (Fig. 2a, Supplementary Fig. 2), which forms dimers whose structures are essentially independent of oxidation state^33–35^. In addition to the wild-type sequence, we produced two disulphide-incompetent variants, C137S and H124K/C137S^46^. Native mass spectra of these constructs at 5 μM revealed pure dimers (oxidised), monomers (H124K/C137S), or mixtures of the two (C137S, reduced) (Fig. 2b). This redox-dependent monomerisation is consistent with the major difference we observed in the oligomeric distributions of the full-length HSP27 (Fig. 1a).

**Fig. 2:**
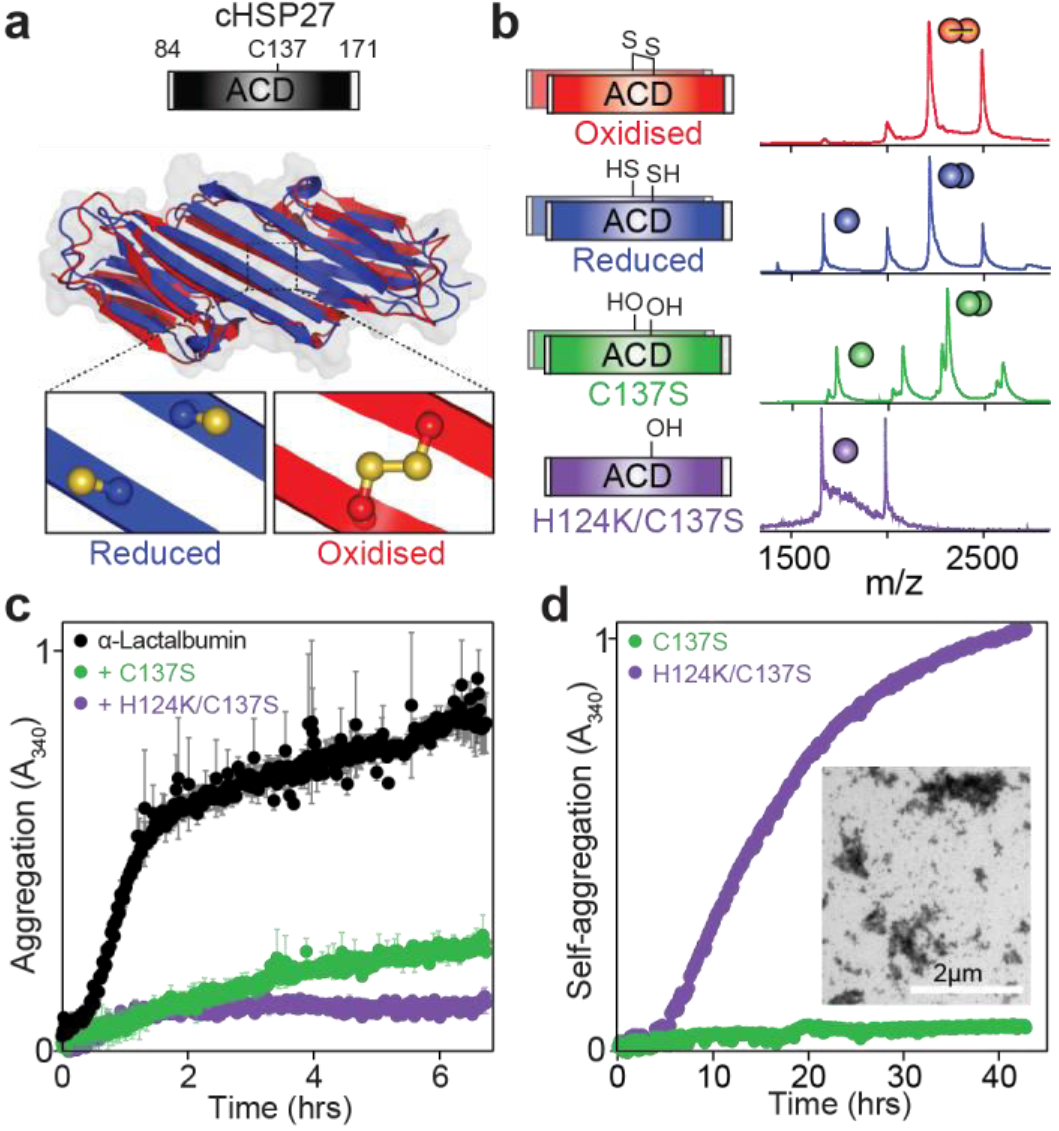
cHSP27 monomers are potent chaperones *in vitro*. (**a**) The central α-crystallin domain (ACD) exists as stable dimers in which residue C137 at the dimer interface can access both reduced (*blue*, PDB 4mjh) and oxidised (*red*, PDB 2n3j) states. (**b**) The cHSP27 variants used in this study together with their native mass spectra at 5 μM, showing the relative population of monomers and dimers present. (**c**) Aggregation of 300 μM dithiothreitol (DTT)-denatured α-lactalbumin (αLac; *black*) in the presence or absence of 70 μM C137S (*green*) or H124K/C137S (*purple*). Under these conditions, C137S is predominantly dimeric whereas H124K/C137S is monomeric. The traces represent the average of three experiments with errors bars indicating ±1 SD. (**d**) The self-aggregation of 700 μM C137S and H124K/C137S was quantified by light scattering at 340 nm at 37 °C over the course of two days. The traces represent the average of six (C137S) or three (H124K/C137S) experiments with error bars indicating ±1 SD. Inset: A representative transmission electron microscopy (TEM) micrograph at 40 hours reveals large, non-fibrillar aggregates of H124K/C137S.

To compare the functional activity of the cHSP27 dimer and monomer, we used the model protein α-lactalbumin (αLac)^34^. At 70 μM, where C137S exists predominantly as a dimer and H124K/C137S a monomer, the double mutant revealed enhanced protection against αLac aggregation (Fig. 2c), further indicating that the monomers are particularly active chaperones. Interestingly, at neutral pH, the H124K/C137S monomer showed a greater propensity than C137S to self-aggregate, forming large amorphous aggregates (Fig. 2d, Supplementary Fig. 3e, 3f) that did not display the Thioflavin-T binding characteristic of amyloid fibrils (Supplementary Fig. 3d). In addition, the thermal stabilities of both reduced cHSP27 and C137S were markedly reduced at low concentrations that favour monomer release (Supplementary Fig. 3). Taken together, these results suggest that, in addition to being more active, the cHSP27 monomer is also kinetically unstable and aggregation prone.

### Dynamics at the dimer interface are redox sensitive

To obtain insight into the structural rearrangements that trigger the enhanced chaperone activity and aggregation propensity of the cHSP27 monomer, we turned to solution-state NMR. 2D ^1^H-^15^N HSQC NMR spectra of oxidised and reduced cHSP27 were recorded at 1 mM, a concentration that favours dimer formation. The spectra were highly similar, a finding that is consistent with the 2.5-Å backbone RMSD between the two forms (Fig. 3a). A quantitative analysis confirmed that both the secondary structure and hydrogen-bonding network (Supplementary Fig. 2) were consistent with published structures^33–35^. Moreover, ^15^N spin relaxation experiments that probe backbone motions revealed that the ps-ns dynamics in cHSP27 were essentially unaltered by changes in redox state (Supplementary Fig. 4). Similarly, we confirmed that C137S effectively mimics the reduced form, as their NMR spectra revealed very similar CSPs to the oxidised form (Fig. 3b), apart for the residues immediately adjacent to the mutation.

Although the structure of the underlying dimer^34,35^ and fast backbone dynamics were redox-independent, when comparing either reduced or C137S to the oxidised form of cHSP27, the signal intensities for residues in the vicinity of the dimer interface were substantially reduced (Supplementary Fig. 2), with elevated ^15^N transverse relaxation rates (Supplementary Fig. 4). These observations demonstrate that reduction of the disulphide bond leads to dynamics on the μs-ms timescale near the dimer interface of cHSP27. Conformational fluctuations between multiple states on this timescale can be characterised using CPMG RD NMR spectroscopy experiments^47,48^, which employ a variable pulse frequency, *ν*_*CPMG*_, to measure effective ^15^N transverse (*R*_2_) relaxation rates^49–51^. The relative populations of the states that are interconverting (*p*_G,_ *p*_E_), their rate of interconversion (*k*_*ex*_), and the chemical shift differences (|*Δω*|) associated with the structural changes can be obtained through quantitative analysis of CPMG RD data.

**Fig. 3:**
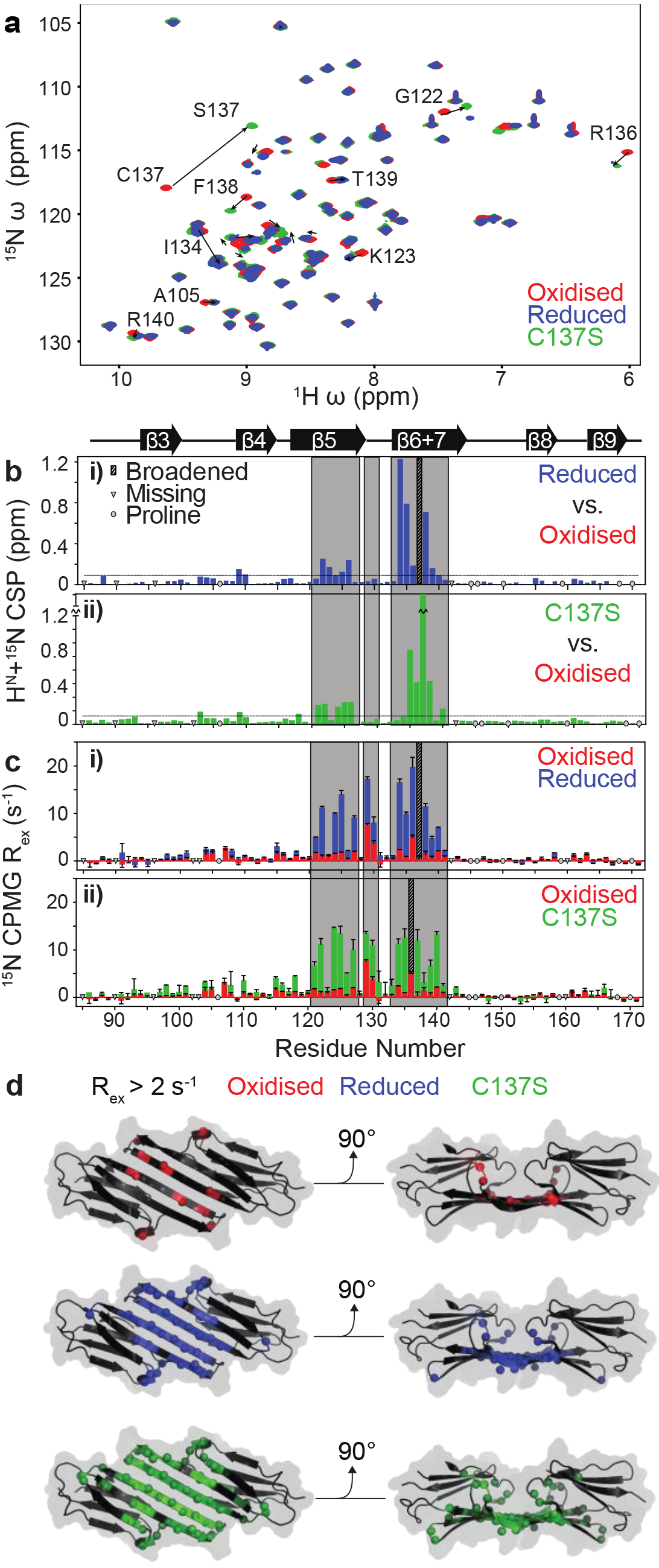
Redox-induced perturbations to the structure and dynamics of cHSP27. (**a**) Overlaid 2D ^1^H-^15^N HSQC spectra of oxidised (*red*), reduced ( *blue*), and C137S (*green*) cHSP27 under identical conditions reveal their similarity. Significant CSPs are indicated with arrows. (**b**) CSPs for individual residues between reduced (**i**) or C137S (**ii**) and the oxidised state reveal differences that localize to the β5 and β6+7 strands. The resonance from C137 was broadened beyond detection in reduced cHSP27 (*cyan*). Proline residues that do not contribute to this spectrum and unassigned residues are indicated. (**c**) Where *R*_*ex*_, approximated here as the difference in *R*_*2,eff*_ at low and high CPMG pulse frequency in ^15^N CPMG experiments, is greater than zero, there is motion on the μs-ms timescale. Residues in the vicinity of L_5,6+7_ show large *R*_*ex*_ values in oxidised, reduced, and C137S cHSP27. The motion extends into the β5 and β6+7 strands for reduced and C137S cHSP27. (**d**) *R*_*ex*_ values > 2 s^−1^ are mapped onto the structure of cHSP27 (PDB 4mjh), indicating that residues with slow dynamics cluster near the dimer interface.

We observed motions on the μs-ms timescale in oxidised, reduced, and C137S cHSP27 (Table S1). H124K/C137S was insufficiently stable to perform the CPMG RD measurements (Fig 2d). In reduced cHSP27 and C137S, the dynamics encompassed β5, L_5,6+7_, and β6+7 (Fig. 3c,d Supplementary Fig. 5). To allow us to characterise the motions in detail we collected an extensive set of CPMG RD data on C137S spanning multiple temperatures (25, 30, and 35°C), concentrations (300 μM and 1 mM), and magnetic field strengths (11.7 and 14.4 T). Quantitative analysis revealed the motion could be interpreted in terms of a global, two-state exchange between the principally populated dimer and a sparsely populated, but partially disordered monomeric state (Fig. 4d, Table S1). From the CPMG RD data, we calculated a *K*_d_ for the C137S monomer-dimer equilibrium of 0.5 μM (Supplementary Figure 5, Table S1), a result qualitatively consistent with results obtained by native MS (Fig. 2 b). Activation and thermodynamic parameters of the monomer-dimer inter-conversion were obtained by analysing the variation in CPMG RD data with temperature. Dissociation of the dimer was endothermic, consistent with a disruption of stabilising interactions, and entropically favoured, consistent with an increase in structural disorder (Table S1). The transition state for dissociation was more disordered than the dimer as evidenced by a positive activation enthalpy and entropy (Supplementary Fig. 5, Table S1).

Similar to reduced cHSP27 and C137S, oxidised cHSP27 exhibited dynamics in L_5,6+7_ (Fig. 4a iii, Supplementary Fig. 5), but these motions were not observed in β5 or β6+7. Detailed analysis revealed that the motions in L_5,6+7_ involve unfolding of the loop, thereby disrupting the intermolecular salt bridge between D129 in L_5,6+7_ and R140 in β6+7 from the adjacent monomer (Fig. 4b), an order-to-disorder transition on the μs-ms timescale that is independent of oxidation state. In addition to the local unfolding of L_5,6+7_, the oxidized form of cHSP27 showed μs-ms motions in the vicinity of residue C137 on a faster timescale, consistent with isomerisation of the disulphide bond^47^ (Supplementary Fig. 5).

### Partially disordered monomers of cHSP27 characterized by RD NMR

**Fig. 4:**
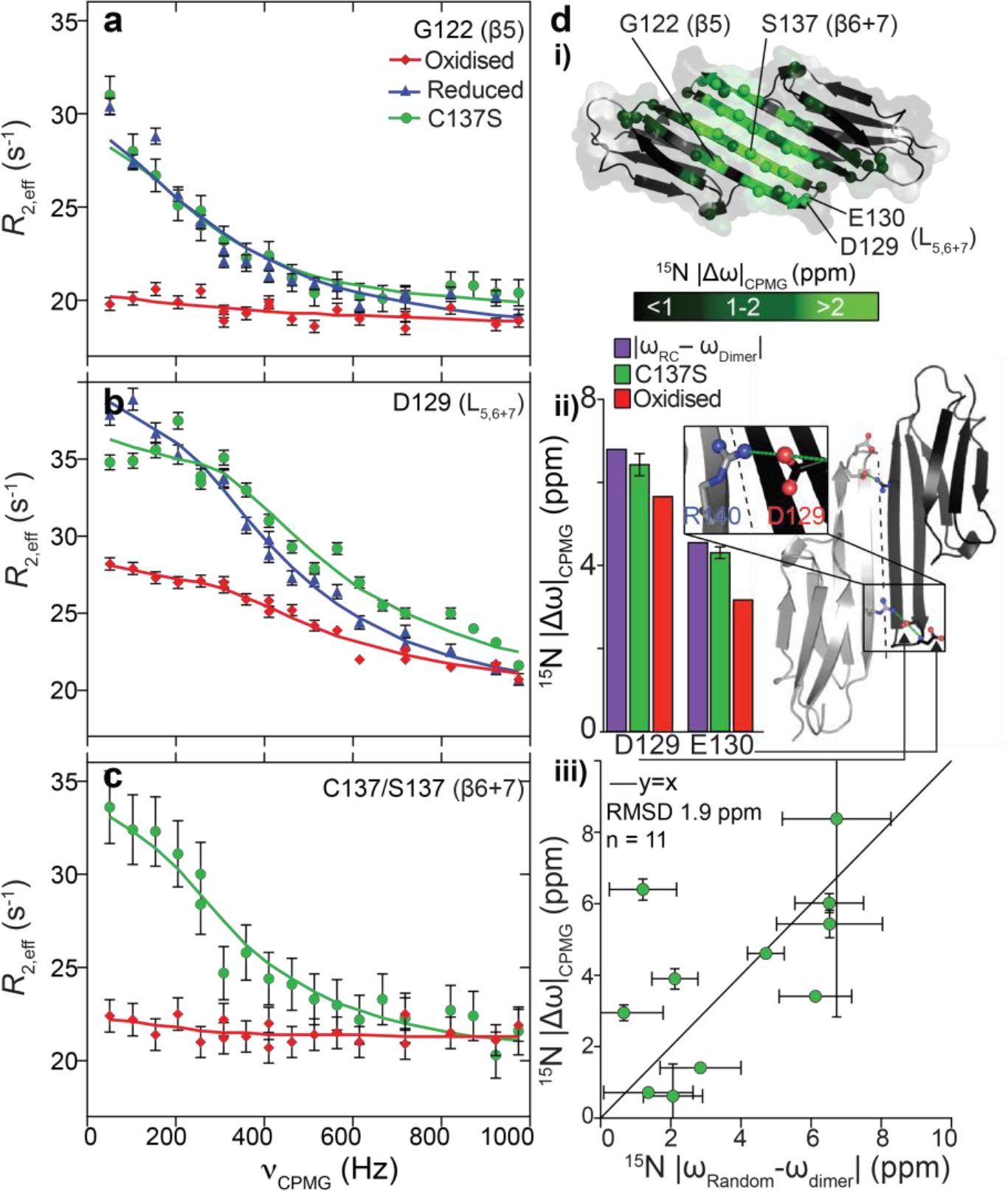
Relaxation dispersion reveals partial unfolding upon transient monomer formation. (**A**) ^15^N CPMG RD experiments quantify μs-ms motions in oxidised (*red*), reduced (*blue*), and C137S (*green*) cHSP27. Fitted curves from a global analysis are shown as solid lines. Significant CPMG RD curves were observed in the β5 strand (**a**), L_5,6+7_ (**b**), and the β6+7 strand (**c**). Redox-independent motions were observed in L_5,6+7_, which arise from unfolding of the loop, whereas only the non-covalent dimers in C137S and reduced cHSP27 show motions throughout β5 and β6+7. (**d i**) CPMG RD-derived ^15^N chemical shift changes in C137S (|Δω|) plotted onto the structure (PDB: 4mjh) revealing significant changes that are localised to L_5,6+7_ and the β5 and β6+7 strands. (**d ii**) The ^15^N |Δω| values in L_5,6+7_ are similar in both C137S and oxidised cHSP27, and correlate with those expected for a transition to a random coil, indicating unfolding of L_5,6+7_ that is independent of oxidation state. In L_5,6+7_, D129 forms an intermolecular salt bridge with R140 from an adjacent subunit, and the amide nitrogen from E130 forms a hydrogen bond with the carbonyl of D129 within the same subunit. (**d iii**) ^15^N |Δω| values in L_5,6+7_ and β6+7 upon monomerisation are compared to the changes expected for random coil formation. The agreement is reasonable, indicating the monomer is substantially disordered in these regions. Outliers are indicated and mainly residue in the end of β6+7.

Given the increased chaperone activity of the monomeric ACD (Fig. 2b), we pursued a structural characterisation of the C137S monomer. The ^15^N chemical shift differences from the CPMG RD experiments (|Δ*ω*|) on C137S report on the structure of the monomeric state. The values that we obtained indicate that residues at the dimer interface (β6+7, L_5,6+7_) adopt ‘random-coil-like’ disordered conformations^48^ (Fig. 4c, Supplementary Fig. 7d). Consistent with this finding, we observed that H124K/C137S displayed characteristics of a partially disordered protein, as evidenced by its 2D ^1^H-^15^N HSQC spectrum (Supplementary Fig. S3a,3b), circular dichroism spectra, and bis-ANS (4,4’-dianilino-1,1’-binaphthyl-5,5’-disulfonic acid) fluorescence. While the majority of the molecule retains its fold upon monomer release, as evidenced by the small |Δ*ω*| values, residues in L_5,6+7_ and β6+7, the region responsible for the dimer interface, become disordered.

### High pressures and acidic conditions enable direct detection of the cHSP27 monomer

While CPMG RD enables a direct characterisation of the HSP27 monomer under near physiological conditions, its low population (ca. 1.5% at 1 mM) renders further high-resolution analysis challenging. We sought to stabilise the monomeric fold. A well-resolved resonance (G116) provided a straightforward marker for distinguishing between the monomeric and dimeric states. Two resonances from this residue were observed in slow exchange at low concentrations for reduced cHSP27 and C137S and the |Δ*ω*| between the two resonances (1 ppm) matched the value obtained from the CPMG analysis. Following the intensities of these resonances allowed us to determine a *K*_d_ for C137S of 0.7 μM, a value consistent with the CPMG analysis (Table S1). Similarly, the concentration dependence of the intensity of the G116 monomer and dimer resonances for H124K/C137S revealed an increase in the *K*_d_ by 3 orders of magnitude to ca. 1.1 mM.

To preserve the monomeric form at sufficiently high concentrations to render it amenable for atomic-resolution characterization by NMR spectroscopy, increased hydrostatic pressure^52^ was employed. NMR spectra of 200 μM C137S were recorded in a baroresistant buffer^53^ at pH 7 as a function of pressure from 1 bar to 2500 bar, revealing a shift in the equilibrium from folded dimer at low pressure to entirely unfolded monomeric C137S at high pressure (Fig. 5a, Supplementary Fig. 6) via an intermediate species that was maximally populated at 1600 bar (Fig. 5b, 5c, Supplementary Fig. 6). The variation in population of dimer, monomer, and unfolded monomer as a function of pressure was explained quantitatively by a three-state linear equilibrium model (Fig. 5b, Table S1). Volumetric changes on application of pressure were obtained, together with the equilibrium constant of monomer unfolding, *K*_*u*_, at 1 bar (Table S1), revealing a free energy difference (Δ_u_*G*) of 5 ± 0.4 kJ mol^−1^ between the monomer and unfolded species at 1 bar and pH 7. The *K*_d_ for dimerisation increased ten-fold at 1600 bar (Fig. 5c, Supplementary Fig. 6). The *K_d_* was further increased by three orders of magnitude by moving from pH 7 to 5 at 1 bar (Fig. 5a, Supplementary Fig. 6, 7). We were able to combine the effects in a phosphate buffer whose pH decreases with pressure, to maximally stabilise the monomeric form (Fig. 5b ii).

While C137S monomer aggregated under acidic conditions at elevated protein concentrations, it remained stable up to 100 μM at pH 4.1 at 1 bar, a finding supported by NMR translational diffusion measurements (Supplementary Fig. 7). Triple resonance spectra for high-resolution analysis of the monomer were acquired under these conditions (Fig. 6a). All observable H^N^, N, Cα, and CO nuclei were assigned (Fig 6a; Supplementary Fig. 7) and, similar to observations by CPMG RD, the largest CSPs fell in L_5,6+7_ and β6+7 (Fig. 6b i, Supplementary Fig. 8). A reasonable correlation was observed (RMSD 1.2 ppm, Supplementary Fig. 7) when the ^15^N chemical shifts from CPMG RD acquired at pH 7 were compared to those measured directly at pH 4.1, indicating that the monomer conformation is similar in both cases. Further confirming their similarity, minor resonances from the monomeric protein could be observed in a sample of C137S at 20 μM at pH 7, whose chemical shifts were close to the values obtained directly under acidic conditions (Fig. 6a, Supplementary Fig. 7).

**Fig. 5:**
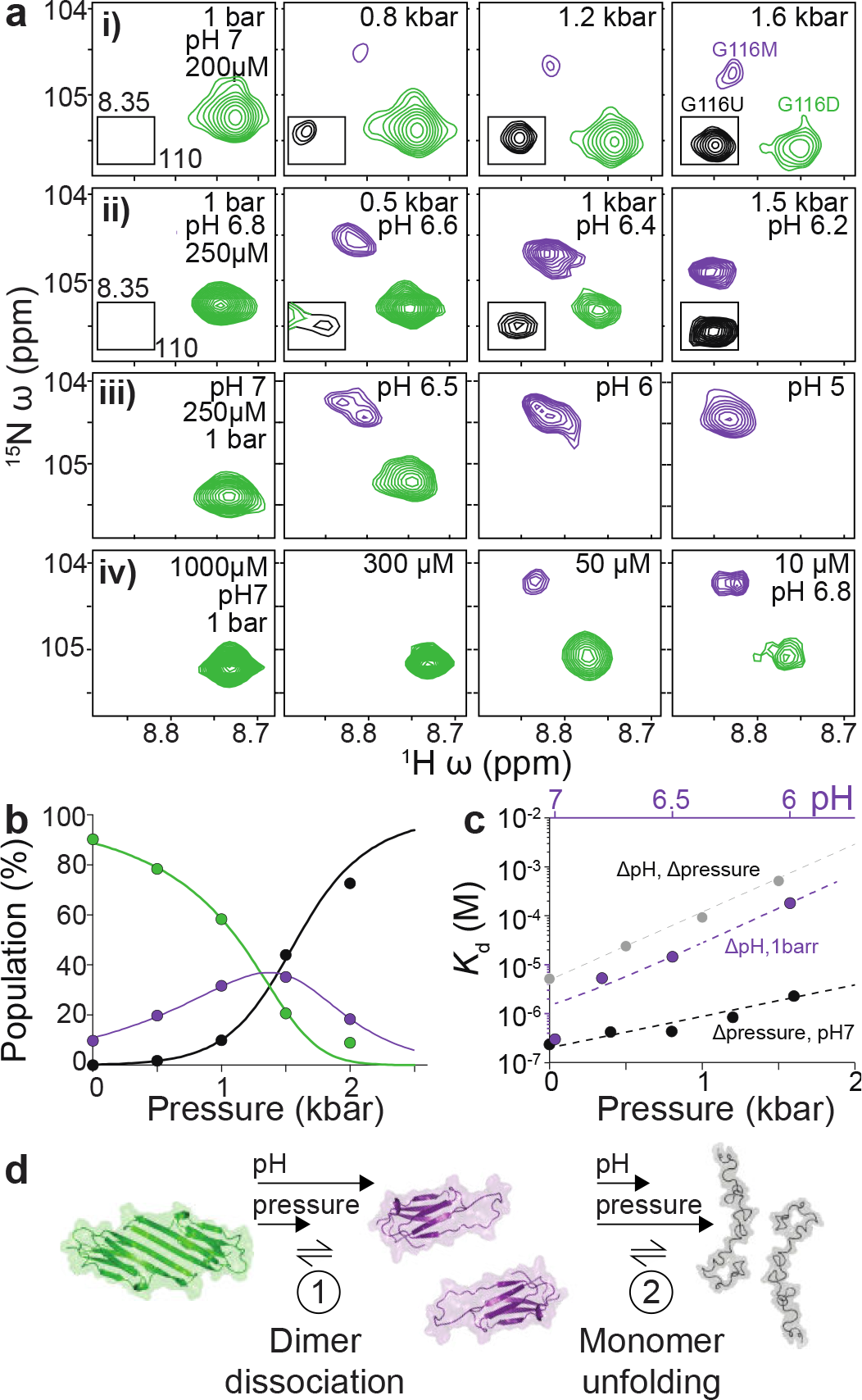
High pressure and low pH stabilize the cHSP27 monomer. (**a**) NMR spectra of C137S, focusing on residue G116 as a function hydrostatic pressure, pH, and concentration. Shown here are (**i**) increasing pressures at pH 7 in a baroresistant buffer (**i**), increasing pressures in phosphate buffer at pH 6.8 at 1 bar, with the pH of phosphate buffer varying with pressure and decreasing by ca. 1 unit over 2 kbar, (**ii**), decreasing pH at 1 bar (**iii**), and decreasing concentration at pH 7 at 1 bar (**iv**). Resonances from the dimer (green), monomer (purple), and unfolded (black) state are readily distinguishable. Decreasing concentration and increasing either pH or pressure favour the monomer over the dimer. At pressures greater than approximately 1.5 kbar, the unfolded form becomes the principally populated state. (**b**) Variations in NMR signal intensities with pressure from four residues that were unambiguously assigned in all three conformations were well explained (solid lines) by the quantitative 3-state equilibrium model shown. (**c**) The *K*_d_ for dimerisation is shown as a function of pressure (*black*), pH (*purple*), and a combination of the two, acquired in phosphate buffer, where the pH of the buffer varies with pressure (grey). (**d**) The three-state mechanism of C137S dimer dissociation, where pH and pressure favour the partially disordered monomer, before ultimate the monomers completely unfold.

### Structural and dynamical characterisation of the partially disordered cHSP27 monomer

To characterise the monomeric state of C137S structurally, we used the observed chemical shifts to determine β-strand formation in the C137S dimer and monomer using the chemical shift^54^ and random coil^55^ indices (CSI, RCI, Fig. 6b ii, Supplementary Fig. 8). This analysis confirmed that, while the disordered L_5,6+7_ spans from Q128 to Q132 in the dimer, it is substantially elongated in the monomer, running from K123 to S137, thereby shortening the β5 and β6+7 strands.

Finally, {^1^H}-^15^N heteronuclear nuclear Overhauser effects (hetNOEs) for the monomer and the dimer were recorded, allowing direct comparison of fast backbone motions on the ps-ns timescale^56^. In the monomer, residues including L_5,6+7_, the C-terminal portion of β5, and the N-terminal portion of β6+7 were highly dynamic (Fig. 6b iii, Supplementary Fig. 8), consistent with the random-coil-like chemical shifts observed in this area by CPMG at pH 7 and directly at pH 4.1. The rigid dimer interface in cHSP27 partially unfolds and becomes highly dynamic in the monomeric state.

**Fig. 6:**
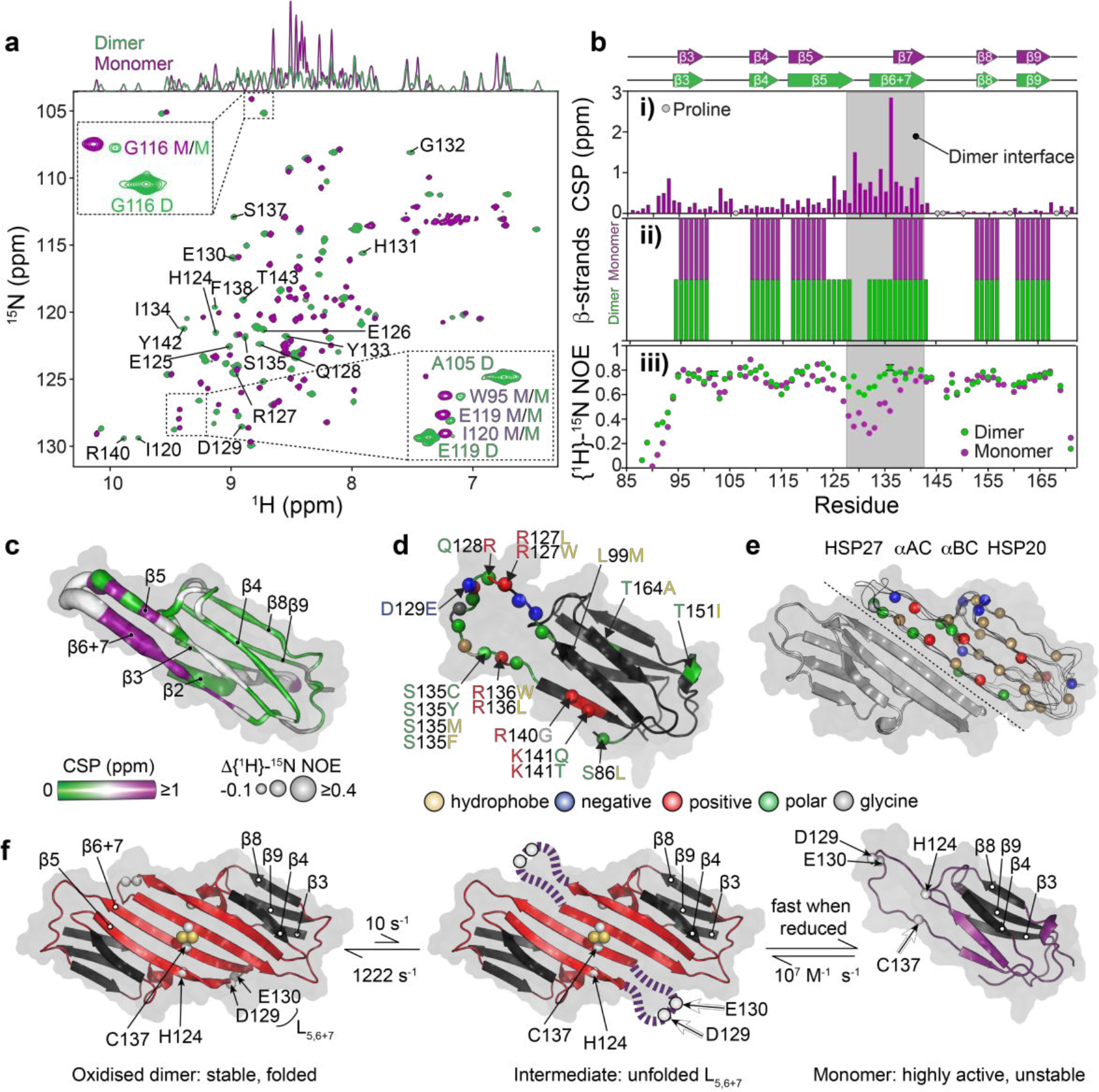
Structural and dynamical characterisation of the partially disordered cHSP27 monomer provides insight into inherited neuropathies. (**a**) *i*. Overlaid NMR spectra of C137S at 20 μM and pH 7 (*green*) where it is predominantly dimeric or pH 4.5 (*purple*) where it is monomeric. The 1D projection onto the ^1^H dimension reveals significant disorder in the monomer. *Insets*: minor resonances observable in the pH 7 sample correspond to the directly observed monomer species at pH 4.5. D/M corresponds to dimer/monomer, respectively. (**b i**) Combined and weighted ^1^H, ^15^N chemical shift perturbations (CSPs) between the C137S dimer at pH 7 and monomer at pH 4.5. (**ii**) β-strands in the C137S monomer (*purple*) and dimer (*green*) as identified by RCI^55^. (**iii**) {^1^H}-^15^N NOEs from the C137S dimer and monomer (Supplementary Fig. 8). (**c**) The ACD monomeric fold from PDB 4mjh with the difference (dimer-monomer) in {^1^H}-^15^N NOE values (tube thickness) and CSPs (colour). (**d**) Inherited mutations that are implicated in the onset of CMT or dHMN disease are indicated in the ACD of HSP27. The mutations cluster to the regions that become solvent exposed upon monomer formation and tend to lower the charge density in the region. (**e**) Overlaid dimer structures of human HSP27 (PDB 4mjh), human αB-crystallin (PDB 4m5s), bovine αA-crystallin (PDB 3l1f), and rat HSP20 (PDB 2wj5) are shown in ribbon format for one subunit of each dimer, and indicate the highly conserved fold of vertebrate ACDs. The second subunit of HSP27 is shown in cartoon format. Highly conserved residues among human sHSPs (HSPB1-HSPB6) are shown as spheres with the same color format as (**f**) The combined results from the CPMG RD and high pressure NMR experiments allow us to propose a hierarchical mechanism for monomer formation. The oxidised, reduced, and C137S forms of cHSP27 exhibit similar dynamics in L_5,6+7_ and form a disordered loop. In the absence of a disulphide bond, this motion in L_5,6+7_ propagates, resulting in the eventual unfolding of the β5 and β6+7 strands in the free monomer.

## Discussion

### HSP27 monomers partially unfold and become highly active

The function and monomerisation of the molecular chaperone HSP27 is regulated by its redox state through an inter-dimer disulphide bond (Fig. 2)^21,23,24^. Here, we determined the structural basis for this regulation. The oligomeric distribution of HSP27 and ps-ns dynamics within the flexible CTR in the full length protein, and the structure and ps-ns dynamics of isolated dimers from excised ACDs were largely invariant to changes in oxidation state. Reduction of full-length HSP27, however, leads to the release of the free monomers (Fig. 1c). Under conditions that favour the free monomer in the context of both the full-length sequences and core domains, we observe enhanced chaperone activity *in vitro* (Fig. 1e). Using a combination of CPMG RD and high pressure NMR, we established that the monomeric protein partly unfolds upon dissociation (Fig. 6), such that the region responsible for the rigid interface in the dimer becomes highly dynamic.

These results suggest that the partly disordered monomer of HSP27 is a particularly active chaperone. This observation is similar to two recent findings, where both the acid-induced unfolding of HdeA and HdeB^57^ and the oxidation dependent unfolding of HSP33^58^ structural forms for aggregation inhibition. Likewise, the homodimeric chaperone CesAB exists in a molten globule-like state with residual helical structure^59^, but undergoes a disorder-to-order transition upon binding to its substrate^60^. More generally, the plastic nature of intrinsically disordered proteins (IDPs) is thought to aid their ability to bind a wide variety of partners^61–64^ via specific, yet transient interactions^65^. It is interesting to speculate that the same mechanism for rapid, promiscuous recognition of binding partners by IDPs is responsible for the heightened activity of partially unfolded chaperones. Interestingly, many of the residues that are unfolded in the HSP27 monomer are charged or polar (Fig. 6d), suggesting that electrostatics may play a role in substrate-recognition, as suggested by bioinformatic analyses^66^. Electrostatically mediated binding has been described for the molecular chaperone Spy^61,62^, but contrasts with the recognition of exposed hydrophobic residues by other molecular chaperones including SecB^67^, trigger factor^68^, DnaK/HSP70^69^, and ClpB/HSP100^70^.

### Mechanism of HSP27 monomer release is hierarchical

Our CPMG RD data inform on a specific mechanism for monomer release. The loop L_5,6+7_ located at the dimer interface of HSP27 undergoes redox-independent motions on the ms timescale, leading it to unfold (Fig. 3, Table S1). When the disulphide bond is present, the local unfolding does not propagate further. However, in the reduced form and C137S variant, the disordering process extends, on the same timescale, into both the end of the β5 and beginning of the β6+7 strands, effectively destabilizing the interface and facilitating monomer release (Fig. 6). Given the conservation of sHSP residues in L_5,6+7_ (Figure 6, Supplementary Fig. 9) and the previously observed millisecond motions in this region of cABC^71,72^, transient unfolding of L_5,6+7_ and the adjacent strands upon monomerisation is likely a common property of mammalian sHSPs.

The process of monomer release and unfolding in HSP27 mirrors the ‘docking and locking’ behaviour observed in a range of protein-protein molecular recognition processes^73^, in which a relatively rapid encounter complex is formed prior to a slower step that provides additional stabilising interactions, which effectively ‘lock’ the complex into place^74^. In the case of HSP27, our data suggest that the β6+7 interface forms relatively quickly, before being ‘locked’ down by the acquisition of structure in L_5,6+7_ and the formation of an inter-molecular salt bridge between D129 and R140 (Fig. 4b).

### Neuropathy-related HSP27 mutations cluster in and near the unfolded region of the monomer

We analysed the positions of 28 mutations in HSP27 that cause either CMT or dHMN (Fig. 6a), including the 17 missense mutations that reside in the ACD^55^. Our structural and dynamical analysis of the cHSP27 monomer reveals that 16 of the 17 mutations in the ACD are located in the disordered region adjacent to or within L_5,6+7_ and the β5 and β6+7 strands. As previously noted^28^, a number of these mutations cluster to the ACD dimer interface. Our NMR data further reveals that the mutations that occur in regions beyond the dimer interface, predominantly fall in regions that are highly disordered in the monomer, suggesting that the behaviour of the monomer is important for understanding the molecular bases of CMT disease and dHMN^75^ perhaps in terms of altered activity, abundance or through causing uncontrolled self-aggregation (Fig. 1g).

While certain mutations decrease chaperone activity, some disease-related HSP27 variants that are more monomeric (e.g. R127W, S135F) and exhibit significantly elevated chaperone activity both *in vitro* and *in vivo*^27,28^. Conversely, disease-related mutants that did not impact monomerisation have shown either no change (T151I) or a decrease (P182L) in activity. These observations suggest that the position of the monomer:dimer equilibrium is an important factor in neuropathies associated with variants of HSP27.

### Disordered monomers are likely a common property of mammalian sHSPs

In light of our findings, we hypothesize that partial unfolding of sHSP ACDs upon monomer release may be a general feature of this class of chaperone. A recent study of cABC showed that the chemical shift changes upon monomer formation are larger for residues located at the dimer interface^71^. We analysed the data and found a strong correlation between ^15^N chemical shift changes in cABC upon monomer formation with those expected for the formation of a random coil (Supplementary Fig. 9). More generally, the dimeric structure and sequence composition of residues at the dimer interface are highly conserved in the mammalian sHSPs HSP27, αB-crystallin, αA-crystallin and HSP20 (Fig. 6e, Supplementary Fig. 9). These results suggest that partial unfolding of monomers upon dissociation is a common property of human sHSPs and that the dimeric building block of sHSP oligomers^36^ is assembled first through partly unfolded monomers. Odd-numbered sHSP oligomers^22^, as encountered in both human sHSPs αB-crystallin and HSP27, will have at least one monomer without a complete dimer interface, indicating that the unstructured monomer can also exist within larger oligomers. Interestingly, for the related αB-crystallin we observed that the dimeric form was more chaperone-active than the monomer, particularly for an amyloidogenic substrate^34^. These differences between the two sHSPs could reflect their contrasting substrate profiles^4^, or multiple binding modes^76^.

### Disordered sHSP monomers as flexible sensors for misfolded proteins

In the context of the isolated ACD, our data suggests that increased disorder in the HSP27 monomer renders it a more potent chaperone *in vitro*. In addition to the partially unfolded ACD, full-length HSP27 contains a disordered 80-residue NTD^77^ and a 28-residue highly flexible CTR^78,79^. The exposed residues from the HSP27 NTD and CTR might also contribute to its mechanism of aggregation inhibition. The inherent plasticity in disordered regions would, in principle, allow for the monomer to sample a wide range of conformational space and thereby facilitate its ability to interact with a diverse set of misfolded target proteins. As the monomer is itself prone to aggregation (Fig. 2d), we speculate that the aggregation-prone contacts in HSP27 are largely responsible for detecting misfolded proteins^80–83^. In the context of the cell, it is would seem undesirable to have high concentrations of aggregation-prone monomers, making it advantageous to store them in oligomers. By holding monomers in this ‘storage’ form, the population of the active but unstable monomeric form is kept both transiently low and highly available^9,41^.

In conclusion, our analysis combining CPMG RD and high-pressure NMR with chaperone and aggregation assays provides the first structural characterisation of the sparsely populated and experimentally elusive monomeric form of HSP27. Although populated at a relatively low abundance in the context of oligomers, the monomeric state is nevertheless particularly active *in vitro*. Part of the region that forms the rigid interface in the dimer unfolds and becomes highly dynamic in the monomer, with the additional structural plasticity in the monomer rendering it a more effective chaperone. However, the monomer is itself more aggregation-prone, whereby off-pathway self-assembly culminates in uncontrolled aggregation. With most sHSPs forming large oligomers, it appears likely that regulation of their self-assembly is a finely tuned proteostatic mechanism inherent to this class of chaperones.

## Materials and Methods

### Protein expression and purification

#### HSP27

All media for growth of cells containing pET(HSP27) plasmids contained 100 μg/mL of Amp. Glycerol stocks of Amp-resistant (Amp^R^) pET(HSP27) plasmids were used to inoculate 5 mL cultures for expression of full-length HSP27. Following growth at 37 °C for 6 hours, the 5 mL cultures were transferred to 100 mL (LB medium) or 50 mL (M9 minimal medium) cultures and grown overnight at 37 °C, which were then used to inoculate 1 L (LB medium) or 500 mL (M9 minimal medium) cultures. When the optical density at 600 nm (OD_600_) of these cultures reached between 0.6 and 0.8 units, IPTG was added to a final concentration of 100 μg/mL and protein expression ensued for 3 hours at 37 °C. Cells were pelleted and frozen at −80 °C until use. For preparation of uniformly-^13^C, ^15^N-labeled ([*U*-^13^C,^15^N]-) HSP27, the M9 minimal medium contained 2 g L^−1^ of [*U*-^13^C]-glucose and 1 g L^−1^ of ^15^NH_4_Cl.

HSP27 was purified in a similar manner as described previously^84^ using anion exchange chromatography (AEX) with HiTrapQ HP columns (GE Healthcare). The column was equilibrated in 20 mM Tris-HCl, 1 mM EDTA, pH 7 (AEX buffer A), after which the lysate was applied. The column was washed with 5% of IEX buffer B (AEX buffer A with 1 M NaCl), and a linear gradient of 5-40% AEX Buffer B followed. HSP27 eluted around 15% AEX Buffer B (150 mM NaCl), after which all HSP27-containing fractions were pooled and concentrated for size exclusion chromatography (SEC). A Superdex S200 26/60 column (GE Healthcare) equilibrated in 20 mM Tris-HCl, 150 mM NaCl, 2 mM EDTA at pH 7 was used to separate HSP27 from the remaining contaminants. HSP27 eluted off the Superdex S200 26/60 near 200 mL, or 0.63 column volumes. After pooling HSP27-containing fractions after SEC, the samples were further purified with a second AEX step using a HiTrap Capto Q ImpRes column (GE Healthcare), with AEX Buffers A and B as described previously. Purified HSP27 was then buffer exchanged using 10K MWCO Amicon spin filters into 30 mM NaH_2_PO_4_, 2 mM EDTA, 2 mM NaN_3_, pH 7.0 with 6% D_2_O (NMR buffer), with an additional 100 mM NaCl for the oxidised state. To study the reduced state, 5 mM BME was added (Fig. 1, S1).

#### cHSP27

DNA encoding the residues 84–171 of HSP27 (cHSP27) was inserted into kanamycin-resistant (Kan^R^) pET28-b plasmids, which contained an N-terminal hexahistidine (His_6_)-tag followed by a TEV protease recognition site^34^. The residual glycine that remains after TEV protease cleavage corresponds to G84 in the HSP27 amino acid sequence. All media for growth of cells containing pET28-b(cHSP27) plasmids contained 30 μg/mL of Kan. cHSP27 cultures were grown as described for full length HSP27. Upon inoculation of the 1 L (LB medium) or 500 mL (M9 minimal medium) cultures, A600 was allowed to reach between 0.6 and 0.8 units, upon which IPTG was added to a final concentration of 100 μg/mL and protein expression continued for 3 hours at 37 °C. Cells were pelleted and frozen at −80 °C until use. For preparation of isotopically labeled cHSP27, cells were grown in H_2_O-based M9 minimal medium as above, supplemented with 4 g L^−1^ of unlabeled glucose, 2 g L^−1^ of [*U*-^13^C]-glucose, 1 g L^−1^ of ^15^NH_4_Cl, or grown in 99.9% D_2_O, respectively, for [*U*-^15^N]-, [*U*-^13^C,^15^N]-, or fractionally deuterated (frac^2^H), [*U*-^15^N]-cHSP27.

cHSP27 was purified in a similar manner as described previously^34^, with initial separation on a Ni^2+^ column that was equilibrated in 20 mM Tris, 150 mM NaCl, 30 mM imidazole, pH 8.0 (IMAC Buffer A). Elution of bound protein from the column was achieved using IMAC Buffer A with 300 mM imidazole, and cHSP27-containing fractions were pooled and incubated with His_6_-tagged TEV protease in IMAC Buffer A at room temperature overnight with 5 mM β-mercaptoethanol (BME) to cleave the His_6_-tag from cHSP27. Cleaved cHSP27 was then separated from His_6_-tagged TEV protease, the cleaved His_6_-tag, and other contaminants using reverse Ni^2+^ affinity separation and then subjected to SEC on a Superdex S75 26/60 column (GE Healthcare), eluting at 170-180 mL, or 0.53-0.56 column volumes. The final SEC buffer contained 20 mM Tris-HCl, 150 mM NaCl, and 5 mM BME at pH 7.0. Using 3K MWCO Amicon spin filters, protein samples were buffer exchanged into and concentrated into NMR buffer for the oxidised state at pH 7 unless otherwise specified. 5 mM BME was added when studying the reduced state (Fig. 2-7, Supplementary Fig. 2-8).

### Site-directed mutagenesis in cHSP27

Primers were designed to encode the single-point mutations H124K and C137S in cHSP27. Site-directed mutagenesis was performed using the QuikChange II Mutagenesis Kit (Agilent), and the correct mutations were verified by DNA sequencing. Protein expression and purification was carried out as above. The final yield of the double mutant, cHSP27(H124K/C137S) (~10 mg/L of *E. coli*) was significantly lower than either WT cHSP27 or C137S (30-40 mg/L of *E. coli*).

### Chaperone activity assays

Porcine heart citrate synthase (CS; Sigma-Aldrich) was buffer exchanged into NMR buffer supplemented with 100 mM NaCl. The aggregation of CS is initiated at high temperatures and does not require the addition of a chemical denaturant (e.g. reducing agents), which enables a direct comparison of the chaperone activity of reduced and oxidised HSP27. The aggregation of 10 μM CS was monitored at 43 °C by following light scattering at 340 nm as a function of time. HSP27 was added to a final concentration of 0.5 μM, a concentration low enough for a substantial concentration of monomers to be expected, either in the absence or presence of 5 mM BME.

Calcium-depleted bovine α-lactalbumin (αLac; Sigma-Aldrich) was dissolved in NMR buffer to a final concentration of 1 mM. Aggregation of αLac is initiated by the addition of reducing agent followed by incubation at high temperatures. The aggregation of αLac (300 μM) at 37 °C was initiated upon the addition of 1 mM DTT, and aggregation was monitored by measuring light scattering at 340 nm as a function of time. cHSP27 variants were added to a final concentration of 70 μM, a concentration where C137S is predominantly a dimer and H104K/C137S a monomer (Fig. 6, Supplementary Fig. 7).

### Native MS

For native MS data acquisition, nanoelectrospray ionization (nESI) experiments were executed according to previously published protocols using instrumental settings optimised to transmit intact protein complexes^85^. A 25 μM sample of HSP27 (purified in the absence of reducing agent) was prepared in 200 mM ammonium acetate (pH 6.9) in the presence and absence of 250 μM DTT (Fig. 1c, Supplementary Fig. 1). cHSP27 samples (5 μM) were prepared in the same buffer with and without 250 μM DTT (Fig. 1d).

### Thermal denaturation of cHSP27

A nanoDSF instrument (NanoTemper) was used to monitor the intrinsic fluorescence of cHSP27 and C137S as a function of temperature. Capillaries contained ~10 μL of cHSP27 that had been prepared in NMR buffer without (oxidised) or with 5 mM BME (reduced). C137S was prepared in NMR buffer without BME. The concentrations of cHSP27 and C137S were 1000, 100, 10, and 1 μM. The initial temperature was 20 °C and was set to increase by 1 °C per minute. Fluorescence readings were recorded at 330 and 350 nm, and the melting temperature (*T*_m_) recorded (Supplementary Fig. 7).

### NMR spectroscopy

#### Backbone and side-chain resonance assignments for dimeric cHSP27

All NMR spectroscopy experiments for resonance assignments of cHSP27 and HSP27 at ambient pressure were recorded on a 14.1 T Varian Inova spectrometer equipped with a 5 mm z-axis gradient triple resonance room temperature probe. 2D ^1^H-^15^N sensitivity-enhanced HSQC spectra^86^ at 14.1 T were typically acquired with ^1^H (^15^N) 513 (64) complex points, spectral widths of 8012 Hz (1800 Hz), maximum acquisition times of 64 ms (35.6 ms), a relaxation delay of 1 s, and four scans per FID for a total acquisition time of 19 minutes. [*U*-^13^C,^15^N]-cHSP27 was prepared at a final concentration of 1 mM in NMR buffer, and [*U*-^13^C,^15^N]-HSP27 was prepared at a final concentration of 2 mM in NMR buffer supplemented with 100 mM NaCl. Standard assignment experiments^87^ were collected, specifically:

HNCO: 766/50/25 complex points and 85.1/39.8/23.6 ms maximum acquisition times for H/C/N dimensions respectively, with 8 scans per FID for a total acquisition time of 12.76 hours.
HN(CA)CO: 1024/40/30 complex points and 85.1/10.6/13.3 ms maximum acquisition times for H/C/N, with 16 scans per FID for a total acquisition time of 23.66 hours
HNCA: 766/40/30 complex points and 85.1/14.8/28.4 ms maximum acquisition times for H/C/N, with 16 scans per transient for a total acquisition time of 24.07 hours.
C(CO)NH: 577/40/20 complex points and 64/4.4/18.9 ms maximum acquisition times for H/C/N, with 64 scans per FID for a total acquisition time of 61.86 hours.
HN(CO)CA: 1024/50/20 complex points sampled at 25% sparsity (H/C/N), 85.1/18.5/18.9 ms maximum acquisition times for H/C/N, with 32 scans per FID for a total acquisition time 8.8 hours.

When NUS was employed, an exponentially-weighted sampling scheme was employed in the indirect dimensions and time-domain data were reconstructed with MddNMR^88^. All 3D NMR spectra at ambient pressure were acquired at 25 °C, processed with NMRPipe^89^, and visualized with Sparky^90^. The resultant ^1^H^N^, ^15^N, ^13^CO, ^13^Cα, and ^13^Cβ chemical shifts were analysed with TALOS-N^91^ and RCI^92^ to respectively estimate the secondary structure and N H order parameters (Supplementary Fig. 2).

#### Backbone resonance assignments for monomeric C137S

Resonance assignments for monomeric C137S were obtained on a 14.1 T Bruker Avance-III spectrometer equipped with a cryogenic probe. A [*U*-^2^H,^13^C,^15^N]-labelled sample at pH 4.1, 2 mM sodium phosphate, 25 °C was prepared at 100 μM. 3D HNCA and HNCO spectra were acquired under these conditions. The HNCO spectrum was acquired with 10% NUS, an inter-scan delay of 1.5 s, 1024/170/140 complex points (H/C/N) with maximum acquisition times of 104/70/85.5 ms, and 4 scans per FID for a total acquisition time of 16 hours. For the HNCA spectrum, 1024/115/84 complex points were acquired for H/C/N with maximum acquisition times of 104/20/58 ms, with 8 scans per FID and an inter-scan delay of 1.4 s for a total acquisition time of 81 hours. A similar set of experiments was recorded on [^2^H,^13^C,^15^N]-labelled C137S dimer at pH 7 in order to compare ^13^C chemical shifts. The NUS spectra were reconstructed with SMILE^93^.

### Indirectly probing hydrogen bonds in oxidised cHSP27

^1^H-^15^N HSQC spectra of [*U*-^15^N]-oxidised cHSP27 were recorded at 293, 295, 298, and 303 K, and the ^1^H chemical shift temperature coefficients (dω^1^H/dT) were calculated (Supplementary Fig. 2). dω^1^H/dT values that are more negative than −4.6 ppb/K are more likely to be solvent exposed and hydrogen bonded to water^94^. Residues with temperature coefficients more positive than −4.6 ppb/K are more likely to be involved in intra- or inter-protein hydrogen bonds. However, it should be noted that residues that are near aromatic rings can yield false positives with values less than −4.6 ppb/K^94^.

To provide an independent NMR dataset that also indirectly probes hydrogen bonds, we buffer exchanged a sample of 1 mM [*U*-^15^N]-oxidised cHSP27 into 99.9% D_2_O and recorded a 2D ^1^H-^15^N HSQC spectrum. The dead time was ~40 min for the entire process and the sample was kept at 4 °C during this time. Intra- or inter-cHSP27 hydrogen bonds involving amide protons were assessed by the presence or absence of signals in the D_2_O ^1^H-^15^N HSQC spectrum (Supplementary Fig. 2).

### ^15^N spin relaxation experiments (T_1_, T_2_, NOE) on cHSP27 and full-length HSP27

Standard pulse sequences to measure ^15^N heteronuclear nuclear Overhauser enhancements (hetNOE), longitudinal (*T*_1_), and transverse (*T*_2_) relaxation times were employed^95^, using 14.1 T and 11.7 T Varian Inova spectrometers equipped with a 5 mm z-axis gradient triple resonance room temperature probes. All experiments at 14.1 T were recorded with 576 (32) complex points, 8992 Hz (1800 Hz) sweep widths, and 64 ms (18 ms) acquisition times, for ^1^H (^15^N). All experiments at 11.7 T were recorded with 512 (32) complex points, 8000 Hz (1519 Hz) sweep widths, and 64 ms (21 ms) acquisition times, for ^1^H (^15^N). *T*_1_ and *T*_2_ values are reported as their respective inverses, *i.e.* rates (*R*_1_ and *R*_2_), for convenience. The spectrometer temperature was calibrated with d_4_-methanol. We recorded ^15^N *R*_1_ (11.7, 14.1 T), *R*_2_ (11.7, 14.1 T), and hetNOE (14.1 T only) values at 25 °C on a 1 mM sample of [*U*-^13^C,^15^N]-labelled oxidised cHSP27 in NMR buffer (Supplementary Fig. 3). Upon completion of these measurements, 5 mM β-mercaptoethanol (BME) was added to the NMR tube, and ^15^N *R*_1_, *R*_2_, and hetNOE values were recorded on reduced cHSP27 at 600 MHz only (Supplementary Fig. 3). Similarly, ^15^N *R*_1_, *R*_2_, and hetNOE experiments at 14.1 T were recorded on oxidised [*U*-^13^C,^15^N]-labelled full-length HSP27, after which 5 mM BME was added to obtain relaxation measurements on the reduced and oxidised species (Supplementary Fig. 2). For data acquired exclusively at 14.1 T, reduced spectral density functions were calculated as described previously^96^. For data acquired at multiple magnetic field strengths (oxidised cHSP27), a model-free analysis was conducted (see below). All data sets mentioned here were processed with NMRPipe^89^, visualized with Sparky^90^, and peak shapes were fit and analysed with FuDA^97^ (http://pound.med.utoronto.ca/~flemming/fuda/). hetNOE experiments on the C137S monomer and dimer were performed on a 14.1 T Bruker Avance-III spectrometer equipped with a cryogenically cooled probe using a previously published pulse sequence^98^.

### Model-free analysis of ^15^N spin relaxation data from oxidised cHSP27

We analysed the ^15^N relaxation data from oxidised cHSP27 at 11.7 T (500 MHz) and 14.1 T (600 MHz) using Lipari-Szabo models^99,100^. Prior to fitting the relaxation data with various Lipari-Szabo models^99,100^, we first determined the rotational diffusion behaviour of cHSP27. Residues with evidence of enhanced ps−ns or μs-ms dynamics, as respectively determined by hetNOE values < 0.65 and R_2_ values > 1 S.D. from the mean, were excluded from the diffusion tensor analysis according to Tjandra *et al.*^101^. After removing dynamic residues, the remaining ^15^N *R*_1_/*R*_2_ ratios were fitted to isotropic (*D*_xx_ = *D*_yy_ = *D*_zz_), axially symmetric (*D*_xx_ = *D*_yy_ ≠ *D*_zz_), and anisotropic (*D*_xx_ ≠ *D*_yy_ ≠ *D*_zz_) diffusion tensors, which were statistically validated by comparing values of χ^2^_red_ and by using F-tests. This analysis indicated that axial symmetry provided the most appropriate diffusion model for cHSP27, which was used to fit ^15^N *R*_2_/*R*_1_ ratios with ModelFree4.15^102^ to determine a global correlation time (τ_c_) of 13.65 ns, which was consistent with a ~20 kDa dimer that exhibits slower diffusion of the axis parallel to the principal component of the diffusion tensor (*D*_‖_/*D*_⊥_ = 1.52). For this analysis, a crystal structure of cHSP27 (PDB 4mjh^34^) was rotated into the frame of the diagonalised diffusion tensor. Values for τ_c_ and D_‖_/D_⊥_ were fixed in the subsequent model-free analysis, in which the N–H bond length was set to 1.02 Å and the ^15^N CSA was set to −160 ppm. Using the five aforementioned values of ^15^N relaxation rates from two magnetic fields, we fit these data using ModelFree4.15^102^ to four models: model 1 (optimized ***S***^2^), model 2 (***S***^2^ and τ_e_), model 3 (***S***^2^ and R_ex_), and model 4 (***S***^2^, τ_e_, and R^ex^). No further benefit was obtained when ***S***^2^_f_ was included in the fitting, and thus was not required for this analysis.

### Analysis of ^15^N CPMG RD data

^15^N CPMG relaxation dispersion (RD) experiments were recorded on 11.7 and 14.1 T NMR spectrometers at 25, 30, and 35 °C using standard pulse sequences^103^. Each data set was recorded with ^1^H (^15^N) 615 (35) complex points (512/29 at 11.7 T), sweep widths of 9615 Hz (1800 Hz) (8000 Hz/1519 Hz), acquisitions times of 64 ms (19 ms) (64 ms/19 ms), and a relaxation delay of 3 seconds. Each experiment was recorded as a pseudo-3D spectrum with the third dimension encoded by the variable delay between pulses in the CPMG pulse train (τ_CPMG_ = 4ν_CPMG_^−1^). For these measurements, frac^2^H, [*U*-^15^N]-labeled samples of cHSP27 (1 mM) or C137S (1 mM and 0.3 mM) were prepared. The experiments contained a fixed constant time of 39 ms for the CPMG period and 20 values of ν_CPMG_ ranging from 54 Hz to 950 Hz. Four scans (1 mM) or eight scans (0.3 mM) per FID were recorded for a total acquisition time of ~10 hours (1 mM) or ~20 hours (0.3 mM). Peak shapes were fit with FuDA^97^ to extract peak intensities, which were then converted into *R*_2,eff_ values using the following relation: *R*_2,eff_(ν_CPMG_) = −*ln*(*I*(ν_CPMG_)/*I*(0) ∗ 1/*T*_relax_), where *I*(ν_CPMG_) is the intensity of a peak at ν_CPMG_, *T*_relax_ is the constant relaxation delay of 39 ms that was absent in the reference spectrum, and *I*(0) is the intensity of a peak in the reference spectrum. Two duplicate ν_CPMG_ points were recorded in each dispersion data set for error analysis, and uncertainties in *R*_2,eff_ were calculated using the standard deviation of peak intensities from such duplicate measurements. From plots of *R*_2,eff_ as a function of ν_CPMG_, *R*_ex_ was estimated by taking the difference of *R*_2_(54 Hz) and *R*_2_(950 Hz). The program CATIA^97^ was used for analysis (Table S1), which accounts for all relevant spin physics including imperfect 180° pulses and differential relaxation between spin states^97^, factors that are not accounted for when using closed form analytical solutions^104^. For each combination of concentration and temperature, data were analysed over multiple field strengths for all residues where *R*_*ex*_ > 2 s^−1^. All residues were assumed to experience a ’two-state’ equilibrium between a majorly populated ‘ground’ state and a sparsely populated ‘excited’ conformational state, and so the same interconversion rates were applied to all residues during analysis. Uncertainties in fitted parameters were estimated by a boot-strapping procedure where synthetic datasets were created by sampling residues with replacement, re-analysing the new dataset and storing the result. The distribution of parameters achieved from 1,000 such operations provided a measure of experimental uncertainty.

Twenty-six residues with *R*_ex_ > 2 s^−1^ at 600 MHz were selected for further analysis (Supplementary Table 2), and their ^15^N CPMG RD data at 14.1 T and 11.7 T were globally fit to a model of two-site chemical exchange using the program CATIA^97^. This analysis enabled extraction of *k*_ex_, *p*_E_, and Δω values (Supplementary Table 1, 2). The global fit from all 26 residues yielded a reduced χ^2^ of 1.46. These residues comprised V101, D107, T110, D115, G116, V117, T121, G122, H124, E125, E126, R127, D129, E130, G132, Y133, I134, S135, R136, S137, F138, T139, R140, K141, and T143. The resultant ^15^N |Δω|_CPMG_ values from C137S were compared to ^15^N chemical shift changes between the folded C137S dimer and random coil conformations (^15^N |Δω|_RC_). The four random coil ^15^N chemical shift databases within the neighbour-corrected intrinsically disordered library (ncIDP^105^) were employed, and the average ^15^N random coil chemical shift and standard deviation was used as the ^15^N |Δω|_RC_, which was then correlated with the measured values of ^15^N |Δω|_CPMG_ (Supplementary Fig. 6).

For oxidised cHSP27, residues D129, E130, R136, C137 and F138 showed significant variation in *R*_*2,eff*_ with ν_CPMG_ (Table S3, S4) (*R*_*ex*_ > 2s^−1^). Analysis assuming each residue had independent exchange rates revealed that D129 and E130 clustered with a *k*_*ex*_ ~1500s^−1^ (Supplementary Table 3), whereas R136, C137 and F138 formed a cluster with *k*_*ex*_ > 3000s^−1^ (Table S4). This distinction was apparent from the raw data (Supplementary Fig. 4) as CPMG RD curves from the latter group showed variation in *R*_*2eff*_ with *ν*_*CPMG*_ at high pulse frequencies, whereas those from the first group were effectively in the fast exchange limit at much low CPMG frequencies. The reduced chi squared value, *χ*^*2*^_red_ of the first group was 1.01, and that of the second group was 1.02.

For reduced cHSP27, CPMG RD data were acquired at 14.1 T exclusively. Residues with *R*_ex_ > 2 s^−1^ were globally fit to a model of two-site chemical exchange, assuming a monomer-dimer exchange event, as above. Twenty-three residues were included in the fit (Supplementary Table 5), comprising V101, D107, T110, D115, G116, V118, T121, G122, K123, H124, E125, E126, R127, D129, E130, Y133, I134, S135, F138, T139, R140, K141, and T143. Residues R136 and C137 were not analysed because these resonances were too broad to provide reliable measurements. The *χ*^*2*^_red_ value from the global fit at one magnetic field strength was 1.45, with *k*_ex_ of ~1400 s^−1^ and *p*_E_ of *~*2%.

### Determination of the dimerization dissociation constant (K_d_) from ^15^N CPMG RD data

The CPMG data were analysed according to a two-state equilibrium scheme where 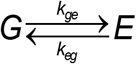. As described in the text, *k*_*eg*_ was found to be concentration dependent, allowing identification of the minor state E to be monomeric, suggesting the equilibrium has the form 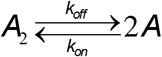. The two-state rate constants derived from CPMG measurements, *k*_*eg*_ and *k*_*ge*_, were converted to *K*_*d*_, *k*_*on*_ and *k*_*off*_ measurements as described in Supplementary Table 1. The *K*_*d*_ values obtained from CPMG analysis from C137S using data acquired at 1 mM and 0.3 mM were highly similar (Supplementary Fig. 4), supporting the identification of the equilibrium to be monomer/dimer exchange.

### Thermal equilibrium and activation analysis of the cHSP27 dimer-to-monomer transition

^15^N CPMG RD data were recorded on 1 mM ^2^H, [*U*-^15^N]-C137S at 25, 30, and 35 °C at both 11.7 T and 14.1 T and analysed globally, assuming the forward and backward rates can be described by the Eyring equation such that 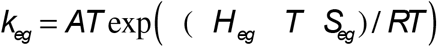 and 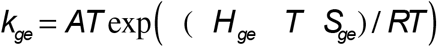. *A* is the reaction frequency, set here to be 3000 s^−1^ K^−1^ and Tis the thermodynamic temperature^106^. Similarly, the equilibrium constant is given by 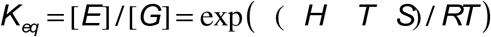 (Supplementary Table 1). Justifying this,locally obtained values of *k*_*eg*_, *k*_*ge*_ and *K*_*eq*_ vary with temperature in a manner consistent with these equations (Supplementary Fig. 4).

### pH-induced dissociation and unfolding of C137S dimers

A 2D ^1^H-^15^N HSQC spectrum was recorded on a sample of [*U*-^15^N]-C137S at 250 μM in NMR buffer at pH 7 at 25 °C. Separate samples were independently prepared in NMR buffer at pH 6.5, 6.0, and 5.0 and NMR spectra were recorded to assess the effect of pH on dimerization. The well-resolved peak from G116 was used to calculate the dimerization *K*_d_ as a function of pH (Supplementary Fig. 5, inset). At pH 5, no dimer was observed, and thus the *K*_d_ at pH 5 (~5 mM) is four orders of magnitude larger than that at pH 7 (0.4 μM). Below pH 6.5, the sample was highly unstable and white precipitant was evident by the end of the NMR experiments (20-40 minutes).

### Analysis of the pressure-induced dissociation and unfolding of C137S dimers

A sample of 200 μM [*U*-^15^N]-C137S was prepared in a baroresistant buffer at pH 7 as described previously^53^ in order to prevent changes in pH with increasing pressure. The buffer contained a mixture of 100 mM Tris-HCl and 100 mM phosphate buffer, both at pH 7, and yields a negligible change in pH between 1 bar and 2500 bar. 2D ^1^H-^15^N HSQC spectra were recorded with ^1^H (^15^N) 1024 (100) complex points, 8417 Hz (2083 Hz) sweep widths, 121.7 ms (48 ms) acquisition times at 14.1 T at 25 °C as a function of hydrostatic pressure between 1 bar to 2500 bar. Manipulation of the hydrostatic pressure was carried out using a commercial ceramic high-pressure NMR cell and an automatic pump system (Daedalus Innovations, Philadelphia, PA). Peak intensities from non-overlapping signals arising from the dimer, monomer, and unfolded species were quantified at each pressure and fit to a model of three-state unfolding (Fig. 5, Supplementary Table 1).

### CD spectroscopy and bis-ANS fluorescence of C137S and cHSP27(H124K/C137S)

Samples for CD spectroscopy were prepared at 20 μM in NMR buffer and placed into cuvettes with a 1 cm path length. Data were recorded using a Jasco Model J720 CD spectrophotometer at wavelengths between 200 and 260 nm. The fluorescent probe bis-ANS (Sigma Aldrich) was dissolved in NMR buffer and added to a final concentration of 10 μM to wells containing a final volume of 100 μL of 40 μM C137S or cHSP27(H124K/C137S) in NMR buffer. Under these conditions, C137S is primarily a dimer and cHSP27(H124K/C137S) is primarily a monomer. Fluorescence at 500 nm was recorded following excitation at 350 nm.

### Aggregation of cHSP27(H124K/C137S)

The aggregation cHSP27(H124K/C137S) was monitored by following the absorbance at 340 nm at 37 °C in NMR buffer. The total volume was 100 μL and the protein concentration was either 200 μM or 800 μM. For comparison, C137S was prepared at 800 μM and subjected to identical treatment. Notably, the sequence-based aggregation propensity predictors Tango^107^ and Zyggregator^108^ do not predict any notable change owning to the choice of mutation. All concentrations had either three or six replicates, and the mean ± SD is reported in Fig. 5 and Supplementary Fig. 7.

### Random Coil Index

To identify β strands in the C137S monomer and dimer, we utilized the software Random Coil Index^55^ and Chemical Shift Index^54^ with default settings. ^1^H^N^, ^13^CO, ^13^Cα, and ^15^N chemical shifts from the C137S dimer (pH 7) and monomer (pH 4.1) were used as input values. The deuterium isotope effect was corrected using established values^109^.

## Acknowledgements

We thank Prof. Heath Ecroyd (University of Wollongong) for the WT HSP27 expression plasmid, Prof. Arthur Laganowsky (Texas A&M University) for assistance in constructing the cHSP27 expression plasmid, Prof. Christina Redfield (University of Oxford) for usage of the CD spectrophotometer, Dr. Errin Johnson (University of Oxford) for assistance with electron microscopy, and Dr. David Staunton (University of Oxford) for assistance with thermal denaturation experiments. TRA acknowledges funding from the NIDDK, the NIH Oxford-Cambridge Scholars Program, and Pembroke College, JLPB thanks the Engineering and Physical Sciences Research Council (EP/J01835X/1) and Biotechnology and Biosciences Research Council (BB/J018082/1), and AJB holds a David Phillips Fellowship from the Biotechnology and Biosciences Research Council (BB/J014346/1). This work was supported in part by the Intramural Research Program of the National Institutes of Diabetes and Digestive and Kidney Diseases and the Intramural Antiviral Target Program of the Office of the Director, NIH.

## Author Contributions

Conceptualization, T.R.A., A.B., J.L.P.B., and A.J.B. Investigation, T.R.A., J.R., H.Y.G., I.P., J.Y., J.L.P.B., A.B., and A.J.B. Writing – Original Draft, T.R.A, A.B., J.L.P.B., and A.J.B. Writing – Reviewing & Editing, T.R.A., J.R., H.Y.G., I.P., J.Y., J.L.P.B., A.B., and A.J.B.

## Additional Materials

Supplementary Figures S1-9, Supplementary Tables S1-5.

## References

1. Kampinga, H. H., de Boer, R. & Beerstra, N. The Big Book on Small Heat Shock Proteins, 3–26: The multicolored world of the human HSPB family. (Springer International Publishing, 2015).

2. Haslbeck, M. & Vierling, E. A first line of stress defense: small heat shock proteins and their function in protein homeostasis. J. Mol. Biol. 427, 1537–48 (2015).

3. Baldwin, A. J., Lioe, H., Robinson, C. V, Kay, L. E. & Benesch, J. L. P. αB-crystallin polydispersity is a consequence of unbiased quaternary dynamics. J. Mol. Biol. 413, 297–309 (2011).

4. Mymrikov, E. V, Daake, M., Richter, B., Haslbeck, M. & Buchner, J. The Chaperone Activity and Substrate Spectrum of Human Small Heat Shock Proteins. J. Biol. Chem. jbc.M116.760413 (2016). doi:10.1074/jbc.M116.760413

5. Bepperling, A. et al. Alternative bacterial two-component small heat shock protein systems. Proc. Natl. Acad. Sci. U. S. A. 109, 20407–12 (2012).

6. Franzmann, T. M., Wühr, M., Richter, K., Walter, S. & Buchner, J. The Activation Mechanism of Hsp26 does not Require Dissociation of the Oligomer. J. Mol. Biol. 350, 1083–1093 (2005).

7. Stengel, F. et al. Quaternary dynamics and plasticity underlie small heat shock protein chaperone function. Proc. Natl. Acad. Sci. U. S. A. 107, 2007–12 (2010).

8. Fleckenstein, T. et al. The Chaperone Activity of the Developmental Small Heat Shock Protein Sip1 Is Regulated by pH-Dependent Conformational Changes. Mol. Cell 58, 1067–1078 (2015).

9. Van Montfort, R., Slingsby, C. & Vierling, E. Structure and function of the small heat shock protein/alpha-crystallin family of molecular chaperones. Adv. Protein Chem. 59, 105–56 (2001).

10. Yu, A. L. et al. Oxidative Stress and TGF-β2 Increase Heat Shock Protein 27 Expression in Human Optic Nerve Head Astrocytes. Investig. Opthalmology Vis. Sci. 49, 5403 (2008).

11. Weindruch, R., Prolla, T. A. & Lee, C.-K. Gene-expression profile of the ageing brain in mice. Nat. Genet. 25, 294–297 (2000).

12. Ciocca, D. R. & Calderwood, S. K. Heat shock proteins in cancer: diagnostic, prognostic, predictive, and treatment implications. Cell Stress Chaperones 10, 86–103 (2005).

13. Outeiro, T. F. et al. Small heat shock proteins protect against α-synuclein-induced toxicity and aggregation. Biochem. Biophys. Res. Commun. 351, 631–638 (2006).

14. Evgrafov, O. V. et al. Mutant small heat-shock protein 27 causes axonal Charcot-Marie-Tooth disease and distal hereditary motor neuropathy. Nat. Genet. 36, 602–6 (2004).

15. Houlden, H. et al. Mutations in the HSP27 (HSPB1) gene cause dominant, recessive, and sporadic distal HMN/CMT type 2. Neurology 71, 1660–8 (2008).

16. Niemann, A. et al. The Gdap1 knockout mouse mechanistically links redox control to Charcot–Marie–Tooth disease. Brain 137, 668–682 (2014).

17. Lin, M. T. & Beal, M. F. Mitochondrial dysfunction and oxidative stress in neurodegenerative diseases. Nature 443, 787–95 (2006).

18. Kirstein, J. et al. Proteotoxic stress and ageing triggers the loss of redox homeostasis across cellular compartments. EMBO J. 34, 2334–49 (2015).

19. Walther, D. M. et al. Widespread Proteome Remodeling and Aggregation in Aging C. elegans. Cell 161, 919–932 (2015).

20. Arrigo, A.-P. et al. Hsp27 consolidates intracellular redox homeostasis by upholding glutathione in its reduced form and by decreasing iron intracellular levels. Antioxid. Redox Signal. 7, 414–22 (2005).

21. Diaz-Latoud, C., Buache, E., Javouhey, E. & Arrigo, A.-P. Substitution of the unique cysteine residue of murine Hsp25 interferes with the protective activity of this stress protein through inhibition of dimer formation. Antioxid. Redox Signal. 7, 436–45 (2005).

22. Zavialov, a et al. The effect of the intersubunit disulfide bond on the structural and functional properties of the small heat shock protein Hsp25. Int. J. Biol. Macromol. 22, 163–73 (1998).

23. Chalova, A. S., Sudnitsyna, M. V, Semenyuk, P. I., Orlov, V. N. & Gusev, N. B. Effect of disulfide crosslinking on thermal transitions and chaperone-like activity of human small heat shock protein HspB1. Cell Stress Chaperones 19, 963–72 (2014).

24. Pasupuleti, N., Gangadhariah, M., Padmanabha, S., Santhoshkumar, P. & Nagaraj, R. H. The role of the cysteine residue in the chaperone and anti-apoptotic functions of human Hsp27. J. Cell. Biochem. 110, 408–19 (2010).

25. Arrigo, A.-P. HSP27: novel regulator of intracellular redox state. IUBMB Life 52, 303–7 (2001).

26. Ye, H. et al. HSPB1 Enhances SIRT2-Mediated G6PD Activation and Promotes Glioma Cell Proliferation. PLoS One 11, e0164285 (2016).

27. Almeida-Souza, L. et al. Increased monomerization of mutant HSPB1 leads to protein hyperactivity in Charcot-Marie-Tooth neuropathy. J. Biol. Chem. 285, 12778–12786 (2010).

28. Almeida-Souza, L. et al. Small heat-shock protein HSPB1 mutants stabilize microtubules in Charcot-Marie-Tooth neuropathy. J. Neurosci. 31, 15320–8 (2011).

29. Hochberg, G. K. A. & Benesch, J. L. P. Dynamical structure of αB-crystallin. Prog. Biophys. Mol. Biol. 115, 11–20 (2014).

30. Aquilina, J. A., Shrestha, S., Morris, A. M. & Ecroyd, H. Structural and functional aspects of hetero-oligomers formed by the small heat shock proteins αB-crystallin and HSP27. J. Biol. Chem. 288, 13602–9 (2013).

31. Jovcevski, B. et al. Phosphomimics Destabilize Hsp27 Oligomeric Assemblies and Enhance Chaperone Activity. Chem. Biol. 22, 186–195 (2015).

32. Rogalla, T. et al. Regulation of Hsp27 oligomerization, chaperone function, and protective activity against oxidative stress/tumor necrosis factor alpha by phosphorylation. J. Biol. Chem. 274, 18947–56 (1999).

33. Baranova, E. V. et al. Three-Dimensional Structure of α-Crystallin Domain Dimers of Human Small Heat Shock Proteins HSPB1 and HSPB6. J. Mol. Biol. 411, 110–122 (2011).

34. Hochberg, G. K. a. et al. The structured core domain of B-crystallin can prevent amyloid fibrillation and associated toxicity. Proc. Natl. Acad. Sci. 111, E1562–E1570 (2014).

35. Rajagopal, P., Liu, Y., Shi, L., Clouser, A. F. & Klevit, R. E. Structure of the α-crystallin domain from the redox-sensitive chaperone, HSPB1. J. Biomol. NMR 63, 223–8 (2015).

36. Mchaourab, H. S., Berengian, A. R. & Koteiche, H. A. Site-Directed Spin-Labeling Study of the Structure and Subunit Interactions along a Conserved Sequence in the α-Crystallin Domain of Heat-Shock Protein 27. Evidence of a Conserved Subunit Interface ^†^. Biochemistry 36, 14627–14634 (1997).

37. Jehle, S. et al. αB-Crystallin: A Hybrid Solid-State/Solution-State NMR Investigation Reveals Structural Aspects of the Heterogeneous Oligomer. J. Mol. Biol. 385, 1481–1497 (2009).

38. Delbecq, S. P., Rosenbaum, J. C. & Klevit, R. E. A Mechanism of Subunit Recruitment in Human Small Heat Shock Protein Oligomers. Biochemistry 54, 4276–4284 (2015).

39. Cox, D., Selig, E., Griffin, M. D. W., Carver, J. A. & Ecroyd, H. Small Heat-shock Proteins Prevent α-Synuclein Aggregation via Transient Interactions and Their Efficacy Is Affected by the Rate of Aggregation. J. Biol. Chem. 291, 22618–22629 (2016).

40. Basha, E., O’Neill, H. & Vierling, E. Small heat shock proteins and α-crystallins: dynamic proteins with flexible functions. Trends Biochem. Sci. 37, 106–17 (2012).

41. Hilton, G. R., Lioe, H., Stengel, F., Baldwin, A. J. & Benesch, J. L. P. Small heat-shock proteins: paramedics of the cell. Top. Curr. Chem. 328, 69–98 (2013).

42. Christodoulou, J. et al. Heteronuclear NMR investigations of dynamic regions of intact Escherichia coli ribosomes. Proc. Natl. Acad. Sci. 101, 10949–10954 (2004).

43. Deckert, A. et al. Structural characterization of the interaction of α-synuclein nascent chains with the ribosomal surface and trigger factor. Proc. Natl. Acad. Sci. 113, 5012–5017 (2016).

44. Cabrita, L. D. et al. A structural ensemble of a ribosome–nascent chain complex during cotranslational protein folding. Nat. Struct. Mol. Biol. 23, 278–285 (2016).

45. Buchner, J., Grallert, H. & Jakob, U. Analysis of chaperone function using citrate synthase as nonnative substrate protein. Methods Enzymol. 290, 323–38 (1998).

46. Clouser, A. F. & Klevit, R. E. pH-dependent structural modulation is conserved in the human small heat shock protein HSBP1. Cell Stress Chaperones 1–7 (2017). doi:10.1007/s12192-017-0783-z

47. Palmer, A. G., Kroenke, C. D. & Loria, J. P. Nuclear magnetic resonance methods for quantifying microsecond-to-millisecond motions in biological macromolecules. Methods Enzymol. 339, 204–238 (2001).

48. Baldwin, A. J. & Kay, L. E. NMR spectroscopy brings invisible protein states into focus. Nat. Chem. Biol. 5, 808–814 (2009).

49. Tollinger, M., Skrynnikov, N. R., Mulder, F. A., Forman-Kay, J. D. & Kay, L. E. Slow dynamics in folded and unfolded states of an SH3 domain. J. Am. Chem. Soc. 123, 11341–52 (2001).

50. Loria, J. P., Rance, M., and & Palmer, A. G. III. A Relaxation-Compensated Carr–Purcell–Meiboom–Gill Sequence for Characterizing Chemical Exchange by NMR Spectroscopy, J. Am. Chem. Soc. 121, 2331–2 (1999).

51. Carver, J. & Richards, R. A general two-site solution for the chemical exchange produced dependence of T2 upon the carr-Purcell pulse separation. J. Magn. Reson. 6, 89–105 (1972).

52. Akasaka, K. Probing conformational fluctuation of proteins by pressure perturbation. Chem. Rev. 106, 1814–35 (2006).

53. Quinlan, R. J. & Reinhart, G. D. Baroresistant buffer mixtures for biochemical analyses. Anal. Biochem. 341, 69–76 (2005).

54. Hafsa, N. E. & Wishart, D. S. CSI 2.0: a significantly improved version of the Chemical Shift Index. J. Biomol. NMR 60, 131–146 (2014).

55. Berjanskii, M. V. & Wishart, D. S. A Simple Method To Predict Protein Flexibility Using Secondary Chemical Shifts. J. Am. Chem. Soc. 127, 14970–1 (2005).

56. Kay, L. E., Torchia, D. A. & Bax, A. Backbone dynamics of proteins as studied by 15N inverse detected heteronuclear NMR spectroscopy: application to staphylococcal nuclease. Biochemistry 28, 8972–8979 (1989).

57. Tapley, T. L. et al. Structural plasticity of an acid-activated chaperone allows promiscuous substrate binding. Proc. Natl. Acad. Sci. U. S. A. 106, 5557–62 (2009).

58. Groitl, B. et al. Protein unfolding as a switch from self-recognition to high-affinity client binding. Nat. Commun. 7, 10357 (2016).

59. Chen, L. et al. Structural Instability Tuning as a Regulatory Mechanism in Protein-Protein Interactions. Mol. Cell 44, 734–744 (2011).

60. Chen, L. et al. Substrate-Activated Conformational Switch on Chaperones Encodes a Targeting Signal in Type III Secretion. Cell Rep. 3, 709–715 (2013).

61. Demarest, S. J. et al. Mutual synergistic folding in recruitment of CBP/p300 by p160 nuclear receptor coactivators. Nature 415, 549–553 (2002).

62. Sugase, K., Dyson, H. J. & Wright, P. E. Mechanism of coupled folding and binding of an intrinsically disordered protein. Nature 447, 1021–1025 (2007).

63. Shammas, S. L., Travis, A. J. & Clarke, J. Allostery within a transcription coactivator is predominantly mediated through dissociation rate constants. Proc. Natl. Acad. Sci. U. S. A. 111, 12055–60 (2014).

64. Ferreon, A. C. M., Ferreon, J. C., Wright, P. E. & Deniz, A. A. Modulation of allostery by protein intrinsic disorder. Nature 498, 390–394 (2013).

65. Wright, P. E. & Dyson, H. J. Intrinsically disordered proteins in cellular signalling and regulation. Nat. Rev. Mol. Cell Biol. 16, 18–29 (2014).

66. Fu, X., Chang, Z., Shi, X., Bu, D. & Wang, C. Multilevel structural characteristics for the natural substrate proteins of bacterial small heat shock proteins. Protein Sci. 23, 229–237 (2014).

67. Huang, C., Rossi, P., Saio, T. & Kalodimos, C. G. Structural basis for the antifolding activity of a molecular chaperone. Nature 537, 202–206 (2016).

68. Saio, T., Guan, X., Rossi, P., Economou, A. & Kalodimos, C. G. Structural Basis for Protein Antiaggregation Activity of the Trigger Factor Chaperone. Science (80-.). 344, 1250494–1250494 (2014).

69. Rosenzweig, R., Sekhar, A., Nagesh, J. & Kay, L. E. Promiscuous binding by Hsp70 results in conformational heterogeneity and fuzzy chaperone-substrate ensembles. Elife 6, (2017).

70. Rosenzweig, R. et al. ClpB N-terminal domain plays a regulatory role in protein disaggregation. Proc. Natl. Acad. Sci. 112, E6872–E6881 (2015).

71. Rajagopal, P. et al. A conserved histidine modulates HSPB5 structure to trigger chaperone activity in response to stress-related acidosis. Elife 4, 1–21 (2015).

72. Mainz, A. et al. Structural and mechanistic implications of metal binding in the small heat-shock protein aB-crystallin. J. Biol. Chem. 287, 1128–1138 (2012).

73. Wright, P. E. & Dyson, H. J. Linking folding and binding. Curr. Opin. Struct. Biol. 19, 31–38 (2009).

74. Schreiber, G. & Fersht, A. R. Rapid, electrostatically assisted association of proteins. Nat. Struct. Biol. 3, 427–431 (1996).

75. Nefedova, V. V., Muranova, L. K., Sudnitsyna, M. V., Ryzhavskaya, A. S. & Gusev, N. B. Small Heat Shock Proteins and Distal Hereditary Neuropathies. Biochem. 80, 1734–1747 (2015).

76. Mainz, A. et al. The chaperone αB-crystallin uses different interfaces to capture an amorphous and an amyloid client. Nat. Struct. Mol. Biol. 22, 898–905 (2015).

77. McDonald, E. T., Bortolus, M., Koteiche, H. A. & Mchaourab, H. S. Sequence, Structure, and Dynamic Determinants of Hsp27 (HspB1) Equilibrium Dissociation Are Encoded by the N-Terminal Domain. Biochemistry 51, 1257–1268 (2012).

78. Carver, J. A., Esposito, G., Schwedersky, G. & Gaestel, M. 1H NMR spectroscopy reveals that mouse Hsp25 has a flexible C-terminal extension of 18 amino acids. FEBS Lett. 369, 305–310 (1995).

79. Alderson, T. R., Benesch, J. L. P. & Baldwin, A. J. Proline isomerization in the C-terminal region of HSP27. Cell Stress Chaperones 22, 639–51 (2017).

80. Carver, J. A. et al. The interaction of the molecular chaperone alpha-crystallin with unfolding alpha-lactalbumin: a structural and kinetic spectroscopic study. J. Mol. Biol. 318, 815–27 (2002).

81. Meehan, S. et al. Characterisation of amyloid fibril formation by small heat-shock chaperone proteins human alphaA-, alphaB- and R120G alphaB-crystallins. J. Mol. Biol. 372, 470–84 (2007).

82. Bardwell, J. C. A. & Jakob, U. Conditional disorder in chaperone action. Trends Biochem. Sci. 37, 517–525 (2012).

83. Jacobs, W. M., Knowles, T. P. J. & Frenkel, D. Oligomers of Heat-Shock Proteins: Structures That Don’t Imply Function. PLoS Comput. Biol. 12, e1004756 (2016).

84. Muranova, L. K., Weeks, S. D., Strelkov, S. V. & Gusev, N. B. Characterization of mutants of human small heat shock protein HspB1 carrying replacements in the N-terminal domain and associated with hereditary motor neuron diseases. PLoS One 10, (2015).

85. Hernández, H. & Robinson, C. V. Determining the stoichiometry and interactions of macromolecular assemblies from mass spectrometry. Nat. Protoc. 2, 715–726 (2007).

86. Kay, L. E., Keifer, P. & Saarinen, T. Pure absorption gradient enhanced heteronuclear single quantum correlation spectroscopy with improved sensitivity. J. Am. Chem. Soc. 114, 10663–10665 (1992).

87. Sattler, M., Schleucher, J. & Griesinger, C. Heteronuclear multidimensional NMR experiments for the structure determination of proteins in solution employing pulsed field gradients. Prog. Nucl. Magn. Reson. Spectrosc. 34, 93–158 (1999).

88. Kazimierczuk, K. & Orekhov, V. Y. Accelerated NMR spectroscopy by using compressed sensing. Angew. Chem. Int. Ed. Engl. 50, 5556–9 (2011).

89. Delaglio, F. et al. NMRPipe: a multidimensional spectral processing system based on UNIX pipes. J. Biomol. NMR 6, 277–93 (1995).

90. Lee, W., Tonelli, M. & Markley, J. L. NMRFAM-SPARKY: enhanced software for biomolecular NMR spectroscopy. Bioinformatics 31, 1325–7 (2015).

91. Shen, Y. & Bax, A. Protein backbone and sidechain torsion angles predicted from NMR chemical shifts using artificial neural networks. J. Biomol. NMR 56, 227–41 (2013).

92. Berjanskii, M. & Wishart, D. S. NMR: prediction of protein flexibility. Nat. Protoc. 1, 683–8 (2006).

93. Ying, J., Delaglio, F., Torchia, D. A. & Bax, A. Sparse multidimensional iterative lineshape-enhanced (SMILE) reconstruction of both non-uniformly sampled and conventional NMR data. J. Biomol. NMR (2016). doi:10.1007/s10858-016-0072-7

94. Baxter, N. J. & Williamson, M. P. Temperature dependence of 1H chemical shifts in proteins. J. Biomol. NMR 9, 359–69 (1997).

95. Palmer, A. G. Dynamic properties of proteins from NMR spectroscopy. Curr. Opin. Biotechnol. 4, 385–391 (1993).

96. Lefevre, J. F., Dayie, K. T., Peng, J. W. & Wagner, G. Internal mobility in the partially folded DNA binding and dimerization domains of GAL4: NMR analysis of the N-H spectral density functions. Biochemistry 35, 2674–2686 (1996).

97. Vallurupalli, P., Hansen, D. F. & Kay, L. E. Structures of invisible, excited protein states by relaxation dispersion NMR spectroscopy. Proc. Natl. Acad. Sci. 105, 11766–11771 (2008).

98. Lakomek, N. A., Ying, J. & Bax, A. Measurement of 15N relaxation rates in perdeuterated proteins by TROSY-based methods. J. Biomol. NMR 53, 209–221 (2012).

99. Clore, G. M. et al. Deviations from the simple two-parameter model-free approach to the interpretation of nitrogen-15 nuclear magnetic relaxation of proteins. J. Am. Chem. Soc. 112, 4989–4991 (1990).

100. Lipari, G. & Szabo, A. Model-free approach to the interpretation of nuclear magnetic resonance relaxation in macromolecules. 1. Theory and range of validity. J. Am. Chem. Soc. 104, 4546–4559 (1982).

101. Tjandra, N., Feller, S. E., Pastor, R. W. & Bax, A. Rotational diffusion anisotropy of human ubiquitin from 15N NMR relaxation. J. Am. Chem. Soc. 117, 12562–12566 (1995).

102. Mandel, A. M., Akke, M. & Palmer, A. G. Backbone dynamics of Escherichia coli ribonuclease HI: correlations with structure and function in an active enzyme. J. Mol. Biol. 246, 144–63 (1995).

103. Sauerwein, A. C. & Hansen, D. F. in 75–132 (Springer US, 2015). doi:10.1007/978-1-4899-7621-5_3

104. Baldwin, A. J. An exact solution for R2,eff in CPMG experiments in the case of two site chemical exchange. J. Magn. Reson. 244, 114–124 (2014).

105. Tamiola, K., Acar, B. & Mulder, F. A. A. Sequence-Specific Random Coil Chemical Shifts of Intrinsically Disordered Proteins. J. Am. Chem. Soc. 132, 18000–18003 (2010).

106. Fersht, A. Enzyme structure and mechanism. (W.H. Freeman, 1985).

107. Linding, R., Schymkowitz, J., Rousseau, F., Diella, F. & Serrano, L. A Comparative Study of the Relationship Between Protein Structure and β-Aggregation in Globular and Intrinsically Disordered Proteins. J. Mol. Biol. 342, 345–353 (2004).

108. Tartaglia, G. G. & Vendruscolo, M. The Zyggregator method for predicting protein aggregation propensities. Chem. Soc. Rev. 37, 1395 (2008).

109. Cavanagh, J., Fairbrother, W. J., Palmer, A. G., Rance, M. & Skelton, N. J. Protein NMR spectroscopy: principles and practice. (Academic Press, 2007).

